# Accelerating protein engineering with fitness landscape modeling and reinforcement learning

**DOI:** 10.1101/2023.11.16.565910

**Authors:** Haoran Sun, Liang He, Pan Deng, Guoqing Liu, Zhiyu Zhao, Yuliang Jiang, Chuan Cao, Fusong Ju, Lijun Wu, Haiguang Liu, Tao Qin, Tie-Yan Liu

**Author notes:** Contributing authors. These authors contributed equally to this work (Haoran Sun, Liang He, Pan Deng, Guoqing Liu).

## Abstract

Protein engineering holds significant promise for designing proteins with customized functions, yet the vast landscape of potential mutations versus limited lab capacity constrains the discovery of optimal sequences. To address this, we present the ***µ***Protein framework, which accelerates protein engineering by combining ***µ***Former, a deep learning model for accurate mutational effect prediction, with ***µ***Search, a reinforcement learning algorithm designed to efficiently navigate the protein fitness landscape using ***µ***Former as an oracle. ***µ***Protein leverages single mutation data to predict optimal sequences with complex, multi-amino acid mutations through its modeling of epistatic interactions and a multi-step search strategy. Except from state-of-the-art performance on benchmark datasets, ***µ***Protein identified high-gain-of-function multi-point mutants for the enzyme ***β***-lactamase, surpassing the highest known activity level, in wet-lab, trained solely on single mutation data. These results demonstrate ***µ***Protein’s capability to discover impactful mutations across vast protein sequence space, offering a robust, efficient approach for protein optimization.

## 1 Introduction

Protein engineering, a cornerstone of biotechnology, focuses on designing proteins with customized functions that advance technology, agriculture, and medicine [1–5]. By optimizing protein sequences, this field holds vast potential for developing biologic drugs, novel enzymes, and other biotechnological innovations. Central to this endeavor is accurately mapping protein sequences to their functions, a process that forms the foundation for identifying sequences with desired traits. This mapping, known as the fitness landscape, is often complex and rugged due to the intricate interactions, or epistasis, between amino acid residues within protein sequences.

High-throughput experimental techniques like Deep Mutational Scanning (DMS)[6] and Multiplex Assays of Variant Effects (MAVEs)[7] have significantly advanced our ability to measure fitness effects for protein variants, particularly by systematically probing single substitutions at every position in a protein. However, these powerful tools face two major limitations. First, they are unable to explore the full combinatorial sequence space, which grows exponentially with the number of mutations. This makes it impractical to experimentally measure high-order mutation combinations, leaving vast regions of the fitness landscape unexplored [4, 8–10]. Second, these methods rely on experimental readout systems that link protein activity to a measurable outcome, such as cell growth, fluorescence, or binding affinity. While effective for certain fitness traits, many protein functions cannot be easily measured using such high-throughput approaches. These constraints further limit the versatility and generality of DMS and MAVEs in addressing diverse protein engineering objectives, underscoring the need for alternative approaches.

Computational models that predict fitness from limited experimental data offer a promising solution, enabling exploration of sequence-function relationships beyond the reach of current experimental techniques [2–5, 10–24]. In this study, we introduce *µ*Protein, a framework that combines two synergistic components: *µ*Former, a deep learning model designed to capture complex epistatic effects accurately, and *µ*Search, a reinforcement learning-based algorithm for efficiently exploring vast sequence spaces using *µ*Former’s predictions. Within this framework, *µ*Former leverages a pairwise masked language model and three specialized scoring modules to model fitness landscapes, which hence enable *µ*Former to handle complex scenarios, such as multiple mutations, indel mutations, and orphan proteins. *µ*Search complements this by framing protein engineering as a decision-making process, using Proximal Policy Optimization (PPO) with exploration noise to navigate the rugged fitness landscape. Together, these components offer a comprehensive and efficient strategy for identifying high-fitness mutations within complex fitness landscapes, thereby expanding the scope of protein engineering.

Applying this pipeline, we successfully engineered a *β*-lactamase enzyme with enhanced activity against cefotaxime, a substrate that has not been extensively explored in protein engineering tasks. By leveraging single mutation data and exploring multi-site, high-fitness mutations, we identified 47 variants with improved activity over the wild type from a pool of 200 tested samples, with *E. coli* harboring certain variants exhibiting growth rates up to 2,000-fold higher than the wild type.

## 2 Results

### 2.1 Overview of *µ*Protein

*µ*Protein is designed to generalize from limited experimental observations and efficiently explore the vast fitness landscapes of proteins (Fig. 1a). The framework combines *µ*Search and *µ*Former. Starting with a wild-type sequence, *µ*Search, the reinforcement learning framework, autonomously navigates the multi-amino acid mutational landscape, guided by *µ*Former, a pre-trained oracle model that scores the mutational effects of sequences proposed by *µ*Search (Fig. 1b).

**Fig. 1:**
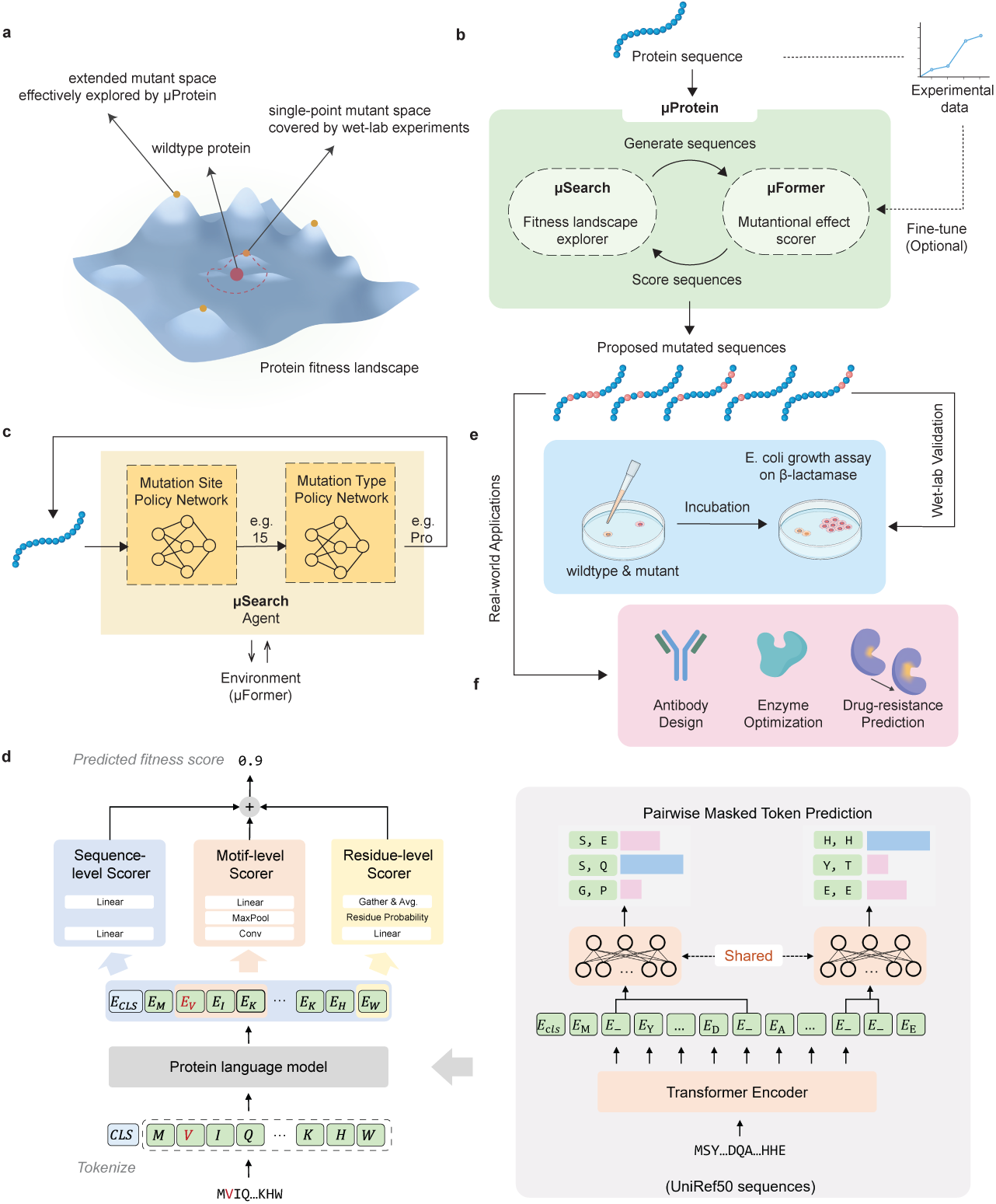
Overview of *µ*Protein. **a)** *µ*Protein efficiently explores the vast protein fitness landscape, generalizing from limited experimental data, typically confined to single-point mutations. **b)** *µ*Protein workflow. **c)** Overview of *µ*Search, a reinforcement-learning algorithm that uses a multi-step search strategy for fitness landscape exploration. **d)** *µ*Former’s two-step mutational effect prediction approach: (1) pre-training of pairwise masked protein language model (PMLM) on a large database of unlabeled proteins to capture dependencies between masked tokens, and (2) integrating three scoring modules (focusing on residual, motif, and sequence-level) for protein fitness prediction. **e)** Wetlab experiment validation of *µ*Protein. **f)** Potential applications of *µ*Protein in real-world scenarios.

*µ*Search frames protein engineering as a Markov Decision Process (MDP), where the state represents the current mutant sequence, the action corresponds to a single amino acid mutation. In each episode of the MDP, *µ*Search sequentially mutates multiple amino acids from the wild-type sequence being engineering. The learning objective of *µ*Search is to optimize the selection policy for mutation sites and types at each step to achieve the desired properties. Using Proximal Policy Optimization (PPO) with Dirichlet noise to avoid local optima, *µ*Search navigates the rugged fitness landscape with both a mutation site policy network and a mutation type policy network, efficiently identifying high-fitness mutations (Fig. 1c. Methods).

*µ*Former, acting as the oracle, provides reward scores for *µ*Search. *µ*Former is built from a self-supervised protein language model combined with three supervised scoring modules (sequence-level, motif-level, and residue-level. Methods). It first undergoes pre-training on over 30 million protein sequences from UniRef50 [25] (Fig. 1d). Using pairwise masked language modeling [26] (Fig. 1d, right), the model learns to predict amino acid identities at masked positions by leveraging both local residue context and pairwise interaction likelihoods within the sequence. During fine-tuning, the full-length sequence embeddings, motif embeddings, and residue probabilities are passed to their respective scoring modules (Fig. 1d, left). The outputs of these modules are then combined to generate the final fitness prediction for the query sequence. The sequence- and motif-level scoring modules have been independently validated as effective tools for modeling protein fitness [22, 27]. Similarly, the residue-level scorer, which evaluates specific amino acid changes at mutated positions, has shown promise in zero-shot fitness prediction tasks [24]. Building on these foundations, we introduce a scoring module that generalizes the residue-level approach. Unlike the traditional residue-level scorer, which focuses exclusively on mutated residues, our design incorporates unmasked sequences during fine-tuning and aggregates probabilities across all residues in the query sequence. The integration of these modules, each focusing on distinct protein features and further enhanced by the pre-trained protein language model, provides a more accurate and comprehensive representation of protein fitness landscapes (Extended Data Fig. 1 and Supplementary Notes), ultimately offering a reliable reward function for *µ*Search.

We validated the *µ*Protein framework through extensive benchmark studies and wet-lab optimization of *β*-lactamase, showcasing its efficacy in pinpointing high-performance mutants in a broader protein mutational space (Fig. 1e). We believe that *µ*Protein holds broad applicability across diverse protein engineering tasks, addressing both industrial and therapeutic challenges, including applications such as antibody design [28], enzyme optimization [29], and the prediction of drug-resistant mutations [30, 31] (Fig. 1f).

### 2.2 *µ*Former accurately predicts fitness scores of proteins

We started by assessing the ability of *µ*Former in fitness landscape modeling and mutational effect prediction. To this end, we benchmarked it against 16 alternative approaches, including MSA-based methods [2, 16–18], language model-based zero-shot methods [5, 24] and learning-based approaches [13–15, 32, 33]. We first evaluated all models on ProteinGym [5], a collection of 78 DMS-derived protein mutational effect datasets spanning a diverse set of protein types, sequence lengths, biological functions, and fitness assays.

Among the tested methods, *µ*Former demonstrates the best overall performance. When analyzing performance on individual datasets, *µ*Former consistently outperforms alternative approaches across the majority of cases (Fig. 2a and Supplementary Data Fig. 1). Moreover, *µ*Former achieves superior results more frequently in terms of Spearman’s rank correlation coefficient (*ρ*) between predicted and observed fitness scores across the tested datasets (Fig. 2b). Notably, for over 50% of the datasets tested, *µ*Former achieves a Spearman’s *ρ* greater than 0.7, and for 6 datasets, it exceeds 0.9, outperforming all other methods. This highlights *µ*Former’s robust capability to deliver highquality predictions across diverse fitness landscapes. As some methods failed on specific datasets due to their suboptimal algorithm design, we calculated both the absolute number and the proportion of high-performing datasets for each model to ensure a fair and comprehensive evaluation.

**Fig. 2:**
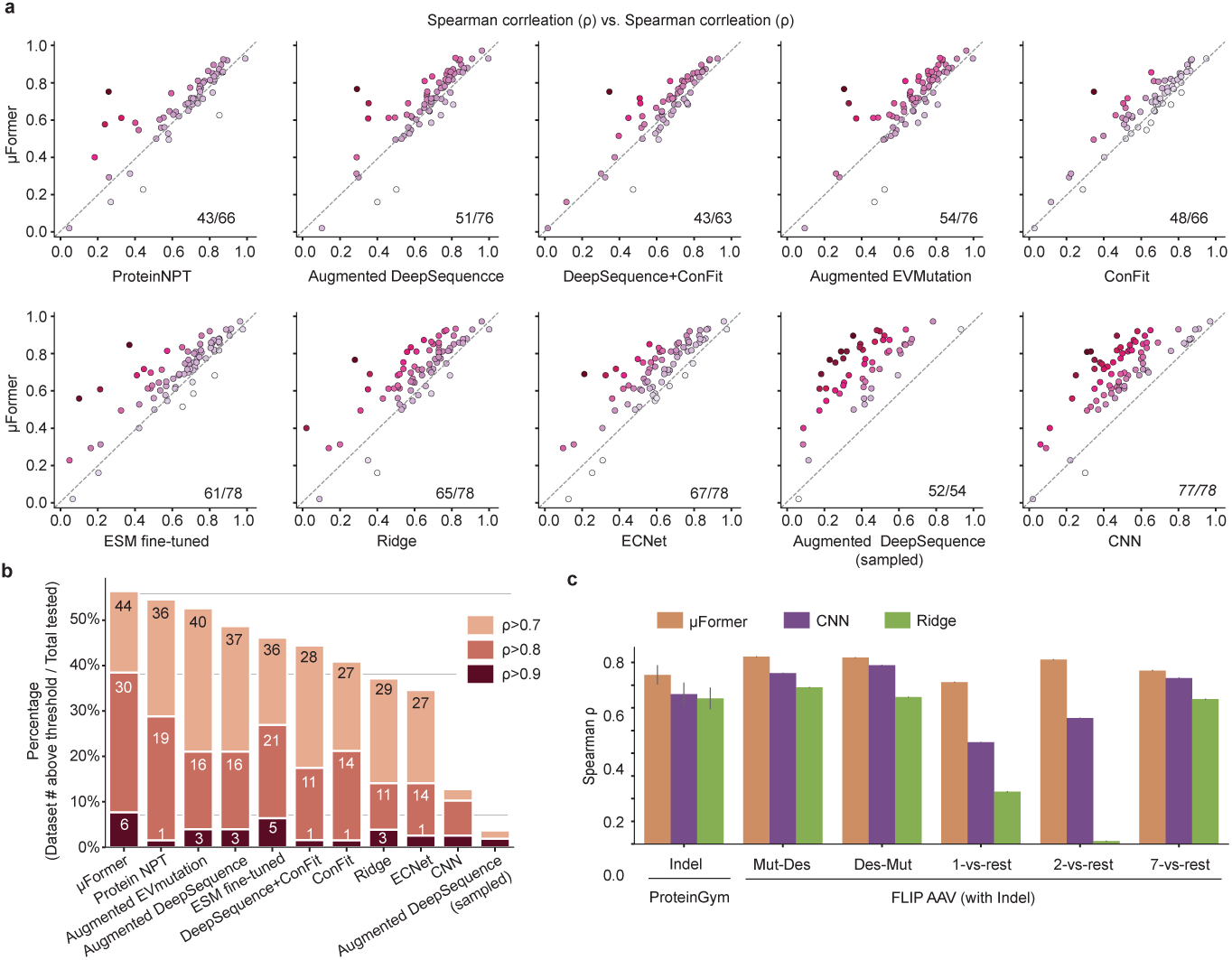
Quantitative comparison of *µ*Former with alternative mutational effect prediction approaches. a) Pairwise comparison between *µ*Former and other approaches. Each data point represents one ProteinGym dataset. Spearman *ρ* is calculated to quantify the performance of corresponding approaches. The lower-right corner displays the number of datasets where *µ*Former outper-formed the corresponding method compared to the total number of tested datasets. The Ridge model refers to the one trained on one-hot embeddings without augmentation. b) The number and percentage of datasets achieving high Spearman correlation for each model. The y-axis represents the percentage of datasets exceeding different Spearman *ρ* thresholds for each model, while the numbers on the bars represent the absolute counts of such datasets. Thresholds of Spearman *ρ* = 0.7, 0.8, and 0.9 were evaluated. Darker bars indicate higher Spearman thresholds. c) *µ*Former outperforms alternative approaches on indel-included benchmark datasets. ProteinGym Indel: cross-protein indelincluded mutational effect dataset collected by Notin et al. [5]. Results from 7 experiments are collectively shown. FLIP AAV: VP1 indel and substitution mutational effect dataset measured and processed by Bryant et al. [34] and Dallago et al. [27] respectively. Results from 10 replications with different random seeds are shown. Mut: naturally occurring mutants. Des: designed sequences. Mut-Des: All natural sequences are assigned to train; all designed sequences are assigned to test. Des-Mut: All designed sequences are assigned to train; all designed sequences are assigned to test. n-vs-rest: Mutants with mutation sites less than or equal to n are assigned to train, the rest of the data with highorder mutations are assigned to test. Center: mean. Error bar: standard error.

Next, we evaluated *µ*Former’s performance on insertion and deletion (indel) prediction. Indels are common mutation types that can lead to drastic changes in protein functions, and designing proteins of different lengths may be expected in specific scenarios. However, indels further complicate the mutational space of proteins, rendering fitness prediction a more challenging task. While MSA-based methods are incapable of scoring indels, approaches of other types may not be able to extend to the scenario of indel prediction due to their designs. Here, we benchmarked *µ*Former on indel tasks against available approaches. Our method showed the best performance on all benchmark datasets with indel mutations (Fig. 2c): ProteinGym Indel, which collected indel-included DMS results from seven assays [5], and FLIP AAV, which split fitness assay data on capsid protein VP1 with varying strategies to probe model generalization [27, 34]. To be noted, the Mut-Des split assigns naturally occurring mutants (Mut) as training dataset and designed sequences (Des) as test dataset. Hence, performances on Mut-Des and the reverse split Des-Mut indicate models’ out-of-distribution prediction ability [27]. *µ*Former achieves a Spearman *ρ* above 0.8 for both settings, signifying its effectiveness as a surrogate oracle for navigating protein fitness landscape and guiding protein sequence design.

We also showed that *µ*Former could generalize to unseen residues, highlighting its data efficiency and generalization ability (Extended Data Fig. 2 and Supplementary Notes)

### 2.3 *µ*Former captures epistatic mutational effect of high-order variants

We next assessed whether *µ*Former could generalize to multi-point mutants, i.e. high-order variants, while being trained exclusively on single-point mutants (single-to-multi setting). We collected nine DMS datasets that include both single-point mutants and high-order mutants [27, 34–41], and trained *µ*Former using only single mutation data for each target protein. To be noted, the training datasets (single mutants) range in size from 362 to 5,468, whereas the test datasets (high-order mutants) are substantially larger (Extended Data Table 1).

A key object in this single-to-multi setting is to assess the model’s ability to predict epistasis, wherein the combined mutational effects at various sites are not simply the sum of their individual effects [8, 42, 43]. Rather than capturing additive effects (linear relationships between single mutations and higher-order mutations), we sought to determine whether models can capture interdependencies between amino acids that are not directly observed in the single-mutant training data. This capability is critical for understanding and modeling complex protein fitness landscapes.

To this end, we calculated epistatic scores for high-order mutants as the discrepancy between the observed mutational effect values and the sum of the constituent single-mutant effect values: 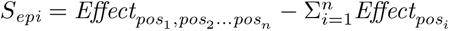 (Fig. 3a). Here, the observed *S_epi_* is defined as the observed multi-mutant effect minus the sum of individual observed single-mutant effects. Similarly, the predicted *S_epi_* is defined as the calculation between the predicted multi-mutant effect and the predicted single-mutant effects generated by each model. Compared to Augmented DeepSequence and ProteinNPT - models that demonstrated strong performance in the previous section - *µ*Former more accurately captures the direction of epistasis, and the epistatic scores predicted by *µ*Former show a stronger correlation with observed *S_epi_* across various proteins (Fig. 3b). Additionally, for high-order mutants exhibiting the top 20% absolute epistatic scores in each protein (Fig. 3a), *µ*Former reduces predictive errors relative to observed values in 8 out of 9 tested proteins compared to the additive model (Fig. 3c). *µ*Former also achieves the highest Spearman’s *ρ* overall (Fig. 3d). These results collectively demonstrate that *µ*Former effectively captures epistatic effects, rather than merely estimating the additive contributions of single mutants present in the training data.

**Fig. 3:**
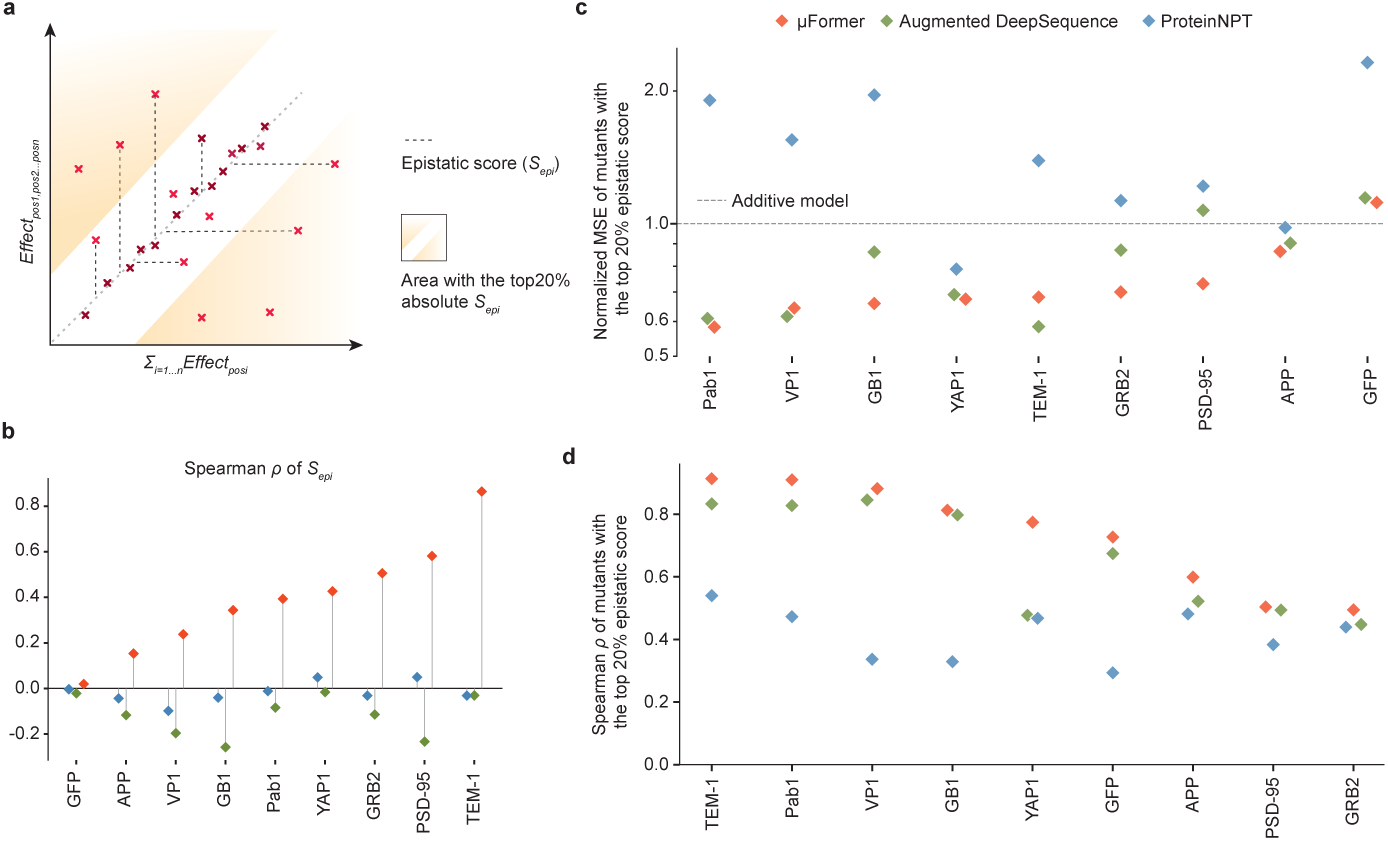
*µ*Former models the epistasis effects in high-order mutants effectively. a) Schematic representation of the calculation of epistatic scores *S_epi_*. High-order mutants with the top 20% absolute *S_epi_* values are highlighted in the shaded regions.b) Spearman *ρ* between predicted and observed *S_epi_* across proteins. c) Normalized mean squared error (MSE) of *µ*Former, ProteinNPT and Augmented DeepSequence for predictions of high-order mutants with the top 20% absolute *S_epi_* values. Compared to the additive model (an epistasis-free estimation), *µ*Former reduces predictive errors in 8 out of 9 tested proteins. d) Spearman *ρ* of *µ*Former, ProteinNPT and Augmented DeepSequence for predictions of high-order mutants with the top 20% absolute *S_epi_* values.

To gain insight into why and how *µ*Former achieves superior performance under the single-to-multi setting, we visualized the embeddings of both single mutants (training data) and high-order mutants (test data) extracted from *µ*Former using t-distributed Stochastic Neighbor Embedding (t-SNE) (Supplementary Data Fig. 2). Interestingly, the embeddings from both pre-trained and fine-tuned models aggregate according to fitness scores, rather than the number of mutations, in the corresponding variants. Thus, the model’s ability to capture the relationship between fitness scores and the corresponding variants may have enabled it to effectively predict the outcomes for high-order mutants, despite not being explicitly trained on them.

### 2.4 *µ*Former effectively identifies high-functioning variants with high-order mutations

The primary goal of protein engineering is to enhance the desired functionality of proteins of interest. While *µ*Former is able to distinguish gain-of-function mutations from loss-of-function mutations accurately (Supplementary Data Fig. 3), we further investigated if the model could predict top-performing mutants effectively.

We calculated the Top-*K* recall scores on curated high-order mutant datasets using the single-to-multi setting for *µ*Former, DeepSequence, and ProteinNPT. This evaluation measures the percentage of top *K* high-functioning high-order mutants correctly identified among the top *K* predictions (hits). In most cases, with *K*=100, *µ*Former provides more than 10 valid hits (Top-100 recall *>* 0.1), and the average Top-100 recall reaches 0.165 (Fig. 4a). When *K* scales up to 500, the recall score further improves and reaches an average value of 0.341 (Fig. 4a). This is a remarkable result, as 100 mutants only represent a small percentage (0.02% to 2%) of tested mutants for each protein. For YAP1, GB1, GRB2 and VP1, a subset of high-order mutants (ultra-high value mutants) exhibits fitness values higher than all single-site mutants of the corresponding protein used for training, which poses an additional challenge to modeling. Encouragingly, *µ*Former makes reasonable predictions for these out-of-distribution variants by detecting ultra-high valued mutants, except the VP1 case (Fig.4a-b). These results together show that *µ*Former can provide efficient guidance for protein optimization.

**Fig. 4:**
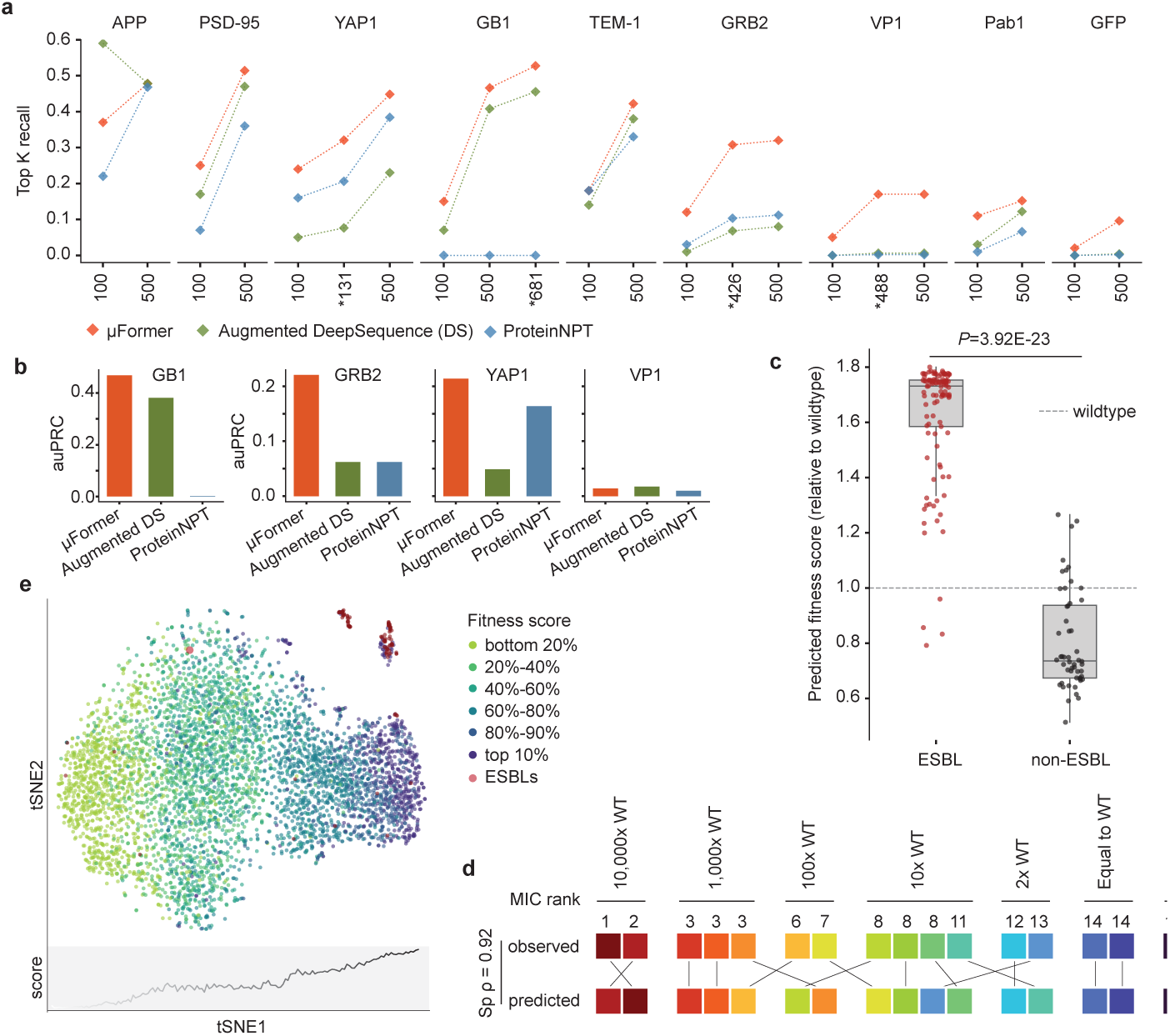
*µ*Former effectively identifies high-functioning variants with multipoint mutations. a) Recall score of *µ*Former, Augmented DeepSequence and ProteinNPT for the Top K high-order mutants of 9 different proteins ranked by fitness values. *K* is equal to 100, 500, or customized values marked with asterisks. The customized values are determined by the number of high-order mutants with fitness values higher than all values in the training data (ultrahigh value mutants). b) The area under precision-recall curve (auPRC) score for *µ*Former and alternate methods in classifying ultra-high value mutants in GB1 (n=681), GRB2 (n=426), YAP1 (n=488), and VP1 (n=131). c) The predicted fitness scores of extended-spectrum *β*-lactamases (ESBLs) are significantly higher than those of non-ESBL clinical variants. The scores are normalized to wildtype (which equals 1). *P* values were computed with the one-sided rank-sum test. n = 157. Center line, median; box limits, upper and lower quartiles; whiskers, 1.5x interquartile range; diamond points, outliers. d) Predicted fitness values from *µ*Former are highly correlated (Spearman *ρ* = 0.92) with the minimum inhibitory concentration (MIC) measured by Weinreich et al [46]. e) t-SNE visualization of TEM-1 mutant embeddings extracted from *µ*Former. Red dots represent ESBLs. Other dots, colored by quantile ranks of experimentally measured fitness values, represent TEM-1 single mutants generated by Stiffler et al [45]. The bottom panel shows the average fitness score along PC1.

To evaluate *µ*Former in practice, we focused on the acquired hydrolysis activity of TEM-1. TEM-1 is a *β*-lactamase that degrades *β*-lactam antibiotics such as ampicillin. Cefotaxime, a *β*-lactam antibiotic discovered nearly 50 years ago, was initially resistant to degradation by TEM-1, making it effective against bacterial pathogens carrying this enzyme. However, TEM-1 variants capable of hydrolyzing cefotaxime rapidly evolved following the drug’s introduction to the market. Among the 222 sequenced clinical variants of TEM-1, 98 are classified as extended-spectrum *β*-lactamases (ESBLs) [44], indicating activity against extended-spectrum cephalosporins with an oxyimino side chain, including cefotaxime. Most ESBLs (91 out of 98) are high-order mutants of TEM-1 (Supplementary Data Fig. 4), we therefore examined whether *µ*Former could prioritize high-order ESBL mutants.

We fine-tuned *µ*Former using a TEM-1 assay performed by Stiffler et al. [45], which measured the fitness effects of approximately 5,000 single-point TEM-1 mutants under cefotaxime selection. Then, we evaluated *µ*Former using a curated dataset comprising 105 ESBLs and 52 confirmed non-ESBLs [44, 46]. *µ*Former prioritizes ESBLs with high scores (Fig. 4c) and demonstrates a significant correlation (Spearman *ρ* = 0.94) with a quantification study on 16 TEM-1 mutants (including 11 high-order mutants) using MIC assay (Fig. 4d) [46]. In contrast, the additive model assigned smaller fitness score gaps between ESBLs and non-ESBLs (Supplementary Data Fig. 5).

To understand how *µ*Former achieves accurate predictions on hyperactive mutants, we extracted mutant embeddings of TEM-1 from *µ*Former and visualized the latent space with t-SNE. On the visualization, an increasing activity of TEM-1 against cefotaxime is clearly observed along the first dimension, with high-functioning variants aggregating on the right (Fig. 4e). Also, when ranked by fitness values, the top 1% (50) TEM-1 mutants from the single mutant DMS study [45] are enriched into 2 isolated clusters on the rightmost, along with 92 out of 105 ESBLs (Fig. 4e). Since the embeddings were extracted from the protein representation layer prior to scoring modules, these results indicate that *µ*Former learns the function of interest of target proteins from sequences, beyond the number of mutations in a mutant sequence.

### 2.5 *µ*Search efficiently navigate fitness landscapes of proteins

*µ*Search is designed to efficiently explore protein fitness landscapes and thus enabling effective protein optimization. To assess the capability of *µ*Search, we first evaluated it using FLEXS [47], an open-source toolbox for benchmarking biological sequence design algorithms. This toolbox provides an approximate fitness model independent of *µ*Former, serving as a noisy oracle to guide diverse exploration algorithms. Additionally, it offers five “ground truth” fitness landscapes spanning DNA, RNA, and proteins [36, 48–51] (Supplementary Notes), enabling comprehensive assessment of exploration algorithms.

We benchmarked *µ*Search against eight alternative exploration algorithms [47, 52–57] across these diverse fitness landscapes. Following the principles of machine-guided directed evolution [58, 59], we employed a multi-round simulated experimental design. In this setup, each algorithm iteratively queried the approximate fitness model, proposed sequences, and updated the model with ground truth oracle-evaluated samples (Supplementary Notes). Specifically, we conducted ten rounds of in silico search using three different random seeds. In each round, each algorithm proposed 100 candidate sequences, constrained by a computational budget of 5,000 local approximate model calls.

Compared to alternative approaches and a Random Selection baseline, *µ*Search demonstrated superior sample efficiency on several protein landscapes (Fig. 5a–c), highlighting its effectiveness in navigating complex protein fitness landscapes. However, we observed that this advantage was less pronounced on nucleic acid-based tasks (Supplementary Data Fig. 6). A possible explanation is that the smaller search spaces in these tasks reduce the difficulty of the optimization problem, resulting in comparable performance across different methods (Supplementary Notes).

**Fig. 5:**
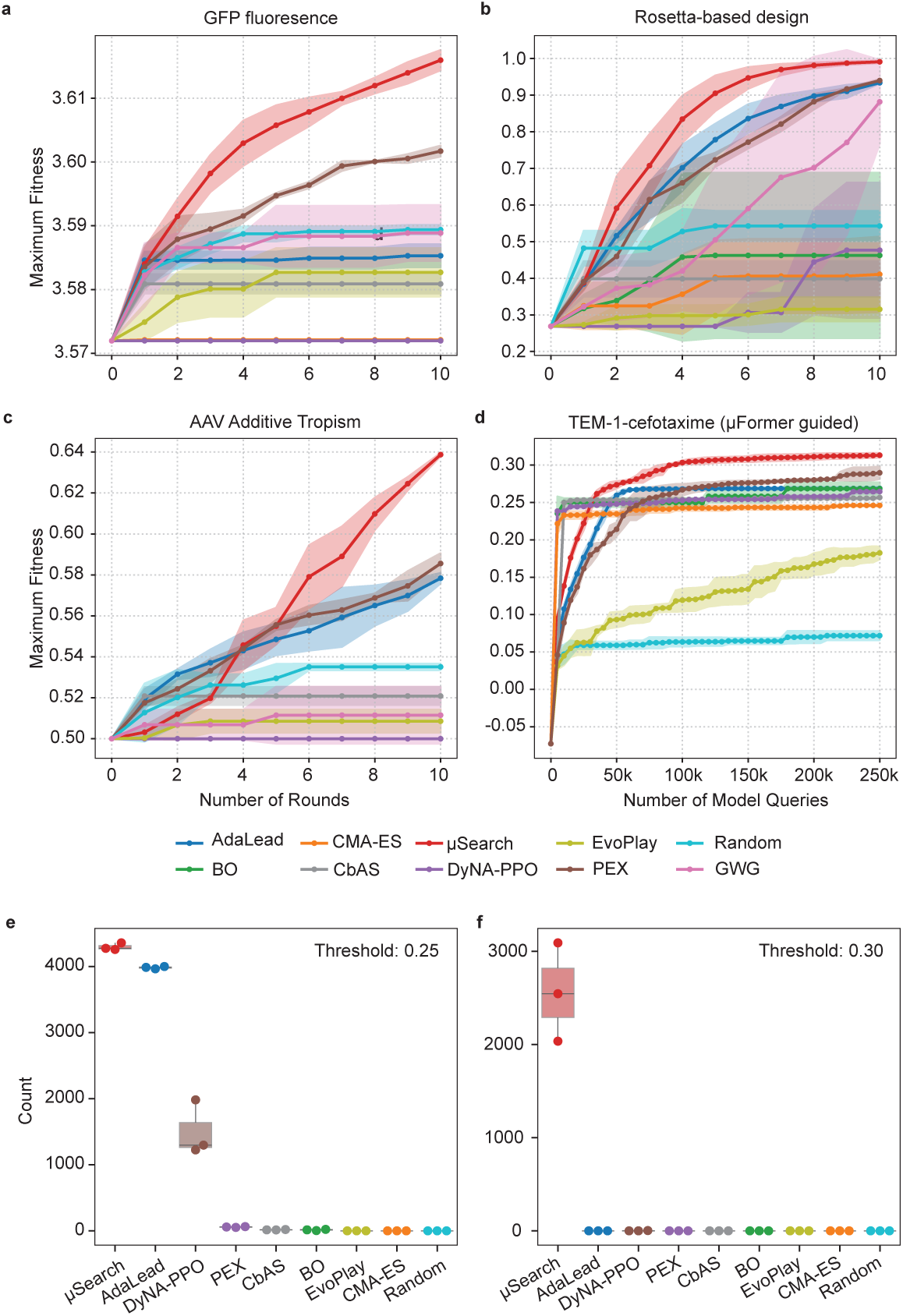
Quantitative comparison of *µ*Search with prevalent fitness landscape exploration algorithms. a-c) Compare *µ*Search with eight leading methods: AdaLead, DyNA-PPO, CbAS, CMA-ES, Bayesian Optimization, PEX, GWG, and EvoPlay, as well as a Random Selection baseline on searching the landscapes of GFP Fluorescence (a), Rosetta-based Design (b), or AAV additive tropism (c) using a multi-round search setting. Maximum fitness: the highest fitness score achieved up to the current round. Center: mean. Error bands: standard deviation. d) Compare *µ*Search with alternative methods using the single-round search setting and *µ*Former as the oracle. A total of 250,000 oracle queries were made for each method. GWG was excluded due to its incompatibility with *µ*Former (It requires joint optimization with an updatable oracle, whereas *µ*Search is decoupled from the oracle, and integrating GWG would require fundamental pipeline changes). Center: mean. Error bands: standard deviation. e-f) The numbers of the mutants that surpass a fitness threshold of 0.25 (e) or 0.3 (f) in the setting for (d). Center line, median; box limits, upper and lower quartiles; whiskers, 1.5x interquartile range; points, outliers. Three

We further evaluated different exploration algorithms using *µ*Former as the oracle model in the TEM-1-cefotaxime system, employing a single-round experimental setting. Here, each algorithm was allowed to make 250,000 oracle queries, without any feedback from the ground truth oracle. *µ*Search identified sequences with higher oracle-predicted fitness scores as the number of model queries increased (Fig. 5d). Remarkably, *µ*Search achieved fitness scores that no other algorithm was able to reach even after 250,000 queries, with around 50,000 queries (Fig. 5d). By the end of 250,000 queries, *µ*Search identified significantly more sequences with high predicted scores compared to other algorithms (Figs. 5e-f). Notably, for sequences with predicted scores exceeding 0.3, *µ*Search identified over 2,000 sequences that are challenging for all other algorithms to detect (Fig. 5f). These results demonstrate that *µ*Search effectively explores complex fitness landscapes and proposes sequences with potentially optimized properties of interest, particularly when paired with *µ*Former.

Lastly, to assess computational efficiency, we compared *µ*Search with random sampling by measuring the number of model queries required to identify mutants exceeding varying fitness thresholds (Supplementary Notes). The results show that *µ*Search achieves the same or higher fitness levels with significantly fewer model evaluations, particularly at higher thresholds, highlighting its superior efficiency in exploration.

### 2.6 Design high-functioning sequences using *µ*Protein

After comprehensive benchmarking on *µ*Former and *µ*Search, we employed the TEM-1-cefotaxime system to assess *µ*Protein’s effectiveness in guiding protein optimization. Using the *µ*Former model fine-tuned on TEM-1 single mutant data (see the previous section) as the reward function, *µ*Search explored millions of possible TEM-1 variants containing 2-3 amino acid changes, aiming to identify those with enhanced activity against cefotaxime (Supplementary Data Fig. 7, Methods).

From the sequences explored through the RL search, we identified over 1,200 variants that exceeded a threshold defined by the median predicted value of known ESBL mutants. From this set, we selected the top 200 variants based on ensemble ranking scores generated by *µ*Former and validated these variants using *E. coli* growth assays (Methods). For comparative analysis, we also included 50 randomly generated variants with 2-3 amino acid changes, as well as 15 variants with 1 to 4 amino acid substitutions spanning A40G, E102K, M180T, and G236S, which have been previously tested in minimum inhibitory concentration (MIC) assays [46],.

*E. coli* carrying 47 distinct RL-designed TEM-1 variants (23.5% of the RL-designed mutants tested) exhibited superior growth on cefotaxime compared to the wild-type strain (Fig. 6a), indicating enhanced enzymatic activity against cefotaxime. In contrast, 12% of randomly generated TEM-1 variants showed enhanced activity. This is consistent with prior observations that TEM-1 mutations often increase activity against cefotaxime [45]. Furthermore, RL-designed variants demonstrated greater activity gains compared to random-generated variants (Fig. 6b). Notably, a double-mutation combination, G236S;T261V, surpassed the activity level of the well-known quadruple mutant A40G;E102K;M180T;G236S, which exhibits significant activity against cefotaxime relative to wild-type TEM-1 [46]. While G236S has been linked to moderate cefotaxime activity gain in previous studies [46], T261I is a mutation that was not previously reported in the ESBL strains. Further validation experiments demonstrate that these novel high-order mutants provide bacteria with a significant growth advantage compared to their constituent single mutations (Supplementary Data Fig. 8).

**Fig. 6:**
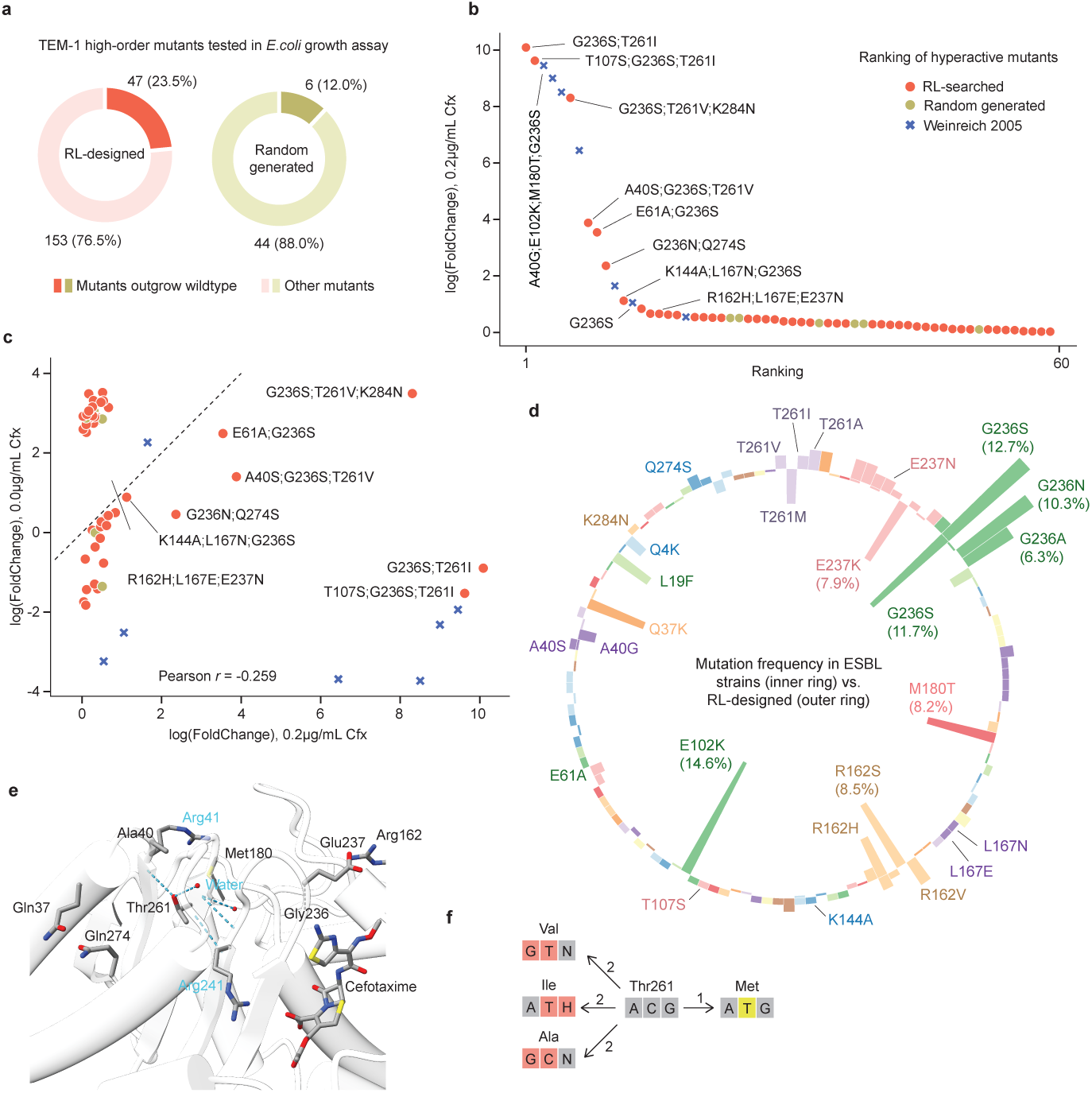
Design high-functioning sequences with *µ*Protein. a) Percentage of TEM-1 high-order mutants that provide a growth advantage to *E. coli* on cefotaxime compared to the wild type, shown for both RL-designed and randomly generated sequences b) Fold change in growth of TEM-1 mutants compared to wild-type *E. coli* in the presence of 0.2 *µ*g/ml cefotaxime, ranked by performance. Only mutants that outgrew the wild-type are shown. c) Comparison of fold change in growth of TEM-1 mutants relative to wild-type *E. coli* under two conditions: 0.2 *µ*g/ml cefotaxime and 0 *µ*g/ml cefotaxime. d) Distribution of mutations in extended-spectrum *β*-lactamases (ESBLs) (inner ring) compared to RL-designed mutants (outer ring). e) The hydrogen bond network involving residue Thr261, with cefotaxime in stick representation. Thr261 forms hydrogen bonds with two water molecules and residues Arg41 and Arg241. f) Illustration depicting nucleotide mutations at position T261 in RL-designed mutants (left) and natural mutants (ESBL strains, right). The numbers 1/2 represent the number of nucleotide mutations required for the amino acid substitution. H denotes A/C/T, and N denotes A/T/C/G.

Comparison of relative growth rates revealed a weak correlation between growth in the presence and absence of cefotaxime for hyperactive mutants (Fig. 6c). Mutants with ultra-high activity tended to exhibit reduced growth in the absence of cefotaxime (bottom-right corner of the plot), consistent with the understanding that beneficial mutations under selective pressure may be detrimental in neutral conditions [60–62]. However, some strains, such as G236S;T261V;K284N, showed growth advantages under both treated and untreated conditions (Fig. 6c).

Notably, the mutation distribution in RL-designed strains differed significantly from that of natural ESBL strains (Fig. 6d). The RL-designed mutants exhibited a more diverse range of mutations. For instance, while the G236S mutation dominates in natural strains, RL-designed mutants frequently featured asparagine and alanine substitutions at G236. Similarly, while T261M is observed in ESBL strains, RL-designed mutants more commonly exhibited isoleucine, valine, and alanine substitutions. All these mutations may alter the hydrogen bond network and, consequently, affect the structure of TEM-1, particularly around arginine at position 241, which is located near the cefotaxime binding pocket and likely influences TEM-1’s activity against the antibiotic (Fig. 6e).

Two factors likely contribute to the observed differences in mutation distribution. First, mutations such as T261A, T261I, and T261V exhibit relatively high activity gains compared to other T261 mutations (Supplementary Data Fig. 9), suggesting that our method favors single mutations with positive effects. Second, natural mutants typically arise from single nucleotide transitions (A-T or C-G mutations), which occur more frequently than transversions *in vivo*. In contrast, RL-designed mutants can explore a broader mutational space, including two-nucleotide mutations (Fig. 6f and Supplementary Data Fig. 10). Therefore, by combining *in silico* fitness landscape modeling with *µ*Search, we are able to design protein variants that may not be achievable through natural evolution or selection. Given that TEM-1 mutants are common pathogenic proteins associated with drug resistance, our findings offer valuable insights into potential future mutations that could arise under prolonged selective pressure.

In conclusion, by leveraging an effective fitness prediction model and an efficient reinforcement learning framework, we successfully navigated the vast mutant space of TEM-1 and identified high-order variants with significantly enhanced activity against cefotaxime. These results represent a significant advancement in exploring and manipulating rugged fitness landscapes.

## 3 Discussion

In this study, we present *µ*Protein, an integrated framework that combines the deep learning model *µ*Former with the reinforcement learning algorithm *µ*Search to model protein fitness landscapes and efficiently explore mutational space. Extensive benchmarking demonstrated *µ*Former’s superior accuracy in predicting fitness landscapes and *µ*Search’s ability to identify optimal variants. We further validated *µ*Protein by optimizing the TEM-1 *beta*-lactamase enzyme. By fine-tuning *µ*Former on TEM-1 single mutant data, we identified several TEM-1 variants highly active against cefotaxime. Notably, *µ*Protein discovered novel high-order mutants, such as G236S;T261V, which exhibited activity levels exceeding those of known high-activity variants. These findings highlight *µ*Protein’s ability to model and search for high-order mutations, addressing the critical challenge of capturing epistatic interactions in protein engineering.

Training data dependency presents a fundamental trade-off in protein engineering, with zero-shot and supervised methods offering distinct advantages. Zero-shot methods are appealing for their ability to generalize without requiring task-specific data, but they often fail to capture desired sequence-function relationships. For proteins like TEM-1 beta-lactamase, where mutational effects can vary significantly across related phenotypes (e.g., resistance to different beta-lactam antibiotics [45], Supplementary Data Fig. 11), supervised training is particularly valuable. Fine-tuning on phenotype-specific data enables the model to achieve higher accuracy in predicting mutational impacts and high-order interactions.

In comparison with alternative methods on extensive benchmarks, *µ*Former demonstrated leading overall performance but did not excel across all tasks. This highlights the fact that no single model currently performs optimally for every protein engineering challenge. Addressing task-specific limitations and enhancing generalizability will be critical for future improvements in computational protein design.

Several potential directions may further enhance *µ*Protein’s capabilities. Integrating 3D structural data may improve its ability to capture spatially proximate epistatic interactions that are not evident from sequence data alone. This would be particularly beneficial for applications like antibody engineering or enzyme optimization, where structural considerations play a key role [63]. Additionally, expanding the training framework to incorporate diverse phenotypes and environmental contexts could enable multitarget fitness predictions within a unified model. For instance, encoding factors such as substrate specificity, temperature, or pH conditions could broaden *µ*Protein’s applicability to more complex protein optimization scenarios.

In this study, we generated approximately 250 TEM-1 mutant strains to assess *µ*Protein’s predictions. While this dataset offered valuable insights, its relatively small size, especially insufficient number of negative samples required for a balanced assessment, limited the fully evaluation on model’s performance. In addition, multi-round experimental feedback, such as iterative cycles of machine learning-guided directed evolution or active learning, could further refine the model and improve its predictive accuracy in future studies.

Overall, *µ*Protein offers a versatile framework for protein engineering by integrating a powerful fitness prediction model with an efficient reinforcement learning strategy. Our study demonstrates its ability to bridge the gap between sequence-based predictions and functional protein optimization. By achieving state-of-the-art performance across a variety of tasks, *µ*Protein provides a valuable tool for designing tailored protein functionalities, opening new possibilities for biotechnological innovation and discovery.

## 4 Methods

### 4.1 Fitness Landscape Modeling via *µ*Former

Our proposed *µ*Former is a deep learning solution for mutation effect prediction, i.e., predicting the fitness score of a mutated protein sequence. Accurate predictions are achieved in two steps: first, we pre-train a masked protein language model (LM) using a large database of unlabeled protein sequences; second, we introduce three scoring modules (each with a small set of new parameters) into the pre-trained protein LM for the final fitness score prediction and train all parameters using a set of mutant protein sequences with measured fitness scores. Figure 1 provides an overview of the prediction model.

In this section, we briefly introduce the masked protein LM and the three scoring modules.

#### 4.1.1 Pairwise Masked Language Model for Proteins

The masked language model (MLM) [19, 64] is a self-supervised learning technique that utilizes the Transformer encoder [65] to learn representations for unlabeled sequences. During training, a random subset of tokens (typically 15%) in the input sequence are masked, and the model is trained to recover the masked tokens, i.e., to minimize the negative log likelihood of masked tokens:

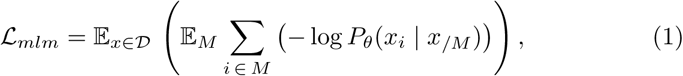

However, protein sequences differ significantly from natural language sentences. Instead of conveying meaning through syntactic and semantic relationships between words, a protein sequence consists of a linear arrangement of amino acids, each with unique physicochemical properties. Together, the amino acids linked sequentially determine a protein’s three-dimensional structure and function. The collective effect of these residues as a whole reflects the sequence’s function, making it essential to learn prHsu et al. protein sequence representations, in order to capture the inter-residue co-variation within the sequences. Conventional masked language models for proteins model masked tokens (i.e., amino acid residues) by conditioning on unmasked tokens only, while processing each token independently. In contrast, we design a pairwise masked language model (PMLM) for protein sequence pre-training, it considers the dependency among masked tokens, taking into account the joint probability of a token pair [26]. One crucial distinction between natural language sentences and protein sequences is that the joint probability cannot be determined by the independent probability of each token. In other words,

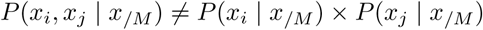

for the *i*-th and *j*-th positions, which represent masked tokens or residues in *M*. This aspect is essential for capturing the co-evolutionary information between elements within a protein sequence. This model is applied to generate a more accurate pre-trained encoder for protein sequence representation.

The loss functions we adopted for protein language model pre-training include the following:

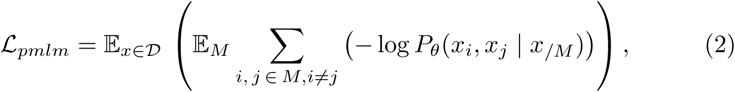

where D represents the set of input sequences, *X* is a sequence in D, *x_/M_* represents the masked version of *x* where the indices of the masked tokens are *M*, *x_i_* is the *i*-th token in the sequence *x*, and *θ* denotes the parameters to be learnt. The final loss function for pre-training is defined as the sum of the masked language modeling loss and the pairwise masked language modeling loss, computed by comparing the ground truths with the predictions from two separate heads:

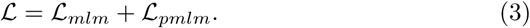

Each Transformer encoder layer consists of two modules: multi-head self-attention (MHA) and position-wise feed-forward (FFN), which are connected through residual connections and layer normalization, as shown below:

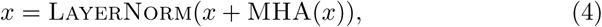

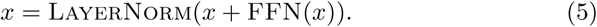

The MLM head and the pairwise MLM head, which predict the masked tokens (i.e., residues in our problem) and the masked token pairs, are both two-layer MLPs that maps the hidden vector at each position to a probability distribution over all possible residues and residue pairs respectively.

In *µ*Former, we pre-train the protein language model using the UR50 dataset, which contains approximately 30 million protein sequences. We adopt the same masking strategy as BERT [19]: for each input sequence, we randomly select 15% of the residues. Of these, 80% are replaced with a special mask token, 10% are substituted with a random amino acid, and the remaining 10% are left unchanged – though the model is still tasked with predicting them and their contributions are included in the loss. This probabilistic behavior enables the model to learn how to refine sequences and identify unfavorable mutations.

#### 4.1.2 Scoring Modules

Motivated by biological insights, we introduce three scoring modules on top of the pre-trained protein LM to predict the fitness score for a protein sequence, which focus on different levels of granularity of the protein sequence, as illustrated in Figure 1:

- The residue-level score *S*_resi_(*x*), which characterizes the likelihood of residues at each position conditioned on the entire sequence [22, 24, 66]. It consists of a multi-layer perceptron (MLP) that processes the hidden representation of each residue to predict amino acid likelihoods. The final score is computed as the average likelihood over all residues.

- The motif-level score *S*_Motif_(*x*), which aims to capture the local sequence information around a residue beyond the residue granularity, considering that motif is widely used in biological sequence modeling for different tasks [10, 64, 67]. It consists of a residual network with two convolutional blocks, followed by a pooling layer and an MLP to generate motif-level scores.

- The sequence-level score *S*_seq_(*x*), is motivated by the observation that the semantics of protein sequences are relevant to their properties/functions on the whole sequence granularity [21, 22, 66, 68, 69]. Like the residue-level scorer, it is an MLP-based model, but it generates predictions using the entire sequence representation as input.

The final fitness score *S*(*x*) is predicted by

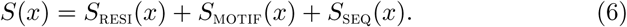

##### Residue-level Scoring Module

This module calculates the fitness score for a mutant-type sequence by averaging the log-likelihood of the residue at each position, as follows:

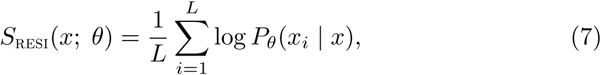

where *L* is the length of the sequence *x*, and *P_θ_*(*x_i_* | *x*) represents the estimated likelihood/probability of the *i*-th residue in the mutant-type sequence *x*, given by the protein LM parameterized by *θ*. Although this score originates from residue-level probabilities, the logits contribute to the final fitness score as a residue-level feature during optimization, where it no longer strictly follows probability constraints. Since the parameters in this component are pre-trained, they also benefit from both the pre-training and fine-tuning processes.

As can be seen, this score only depends on the mutant-type sequence and does not explicitly rely on its wild-type sequence. Notably, previous models [21,24] often use the term 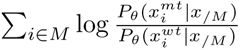 for mutational effect prediction, where *M* is the set of mutated positions, and 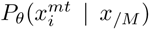 and 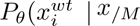) are the estimated probabilities for the *i*-th residues of the mutant-type and wild-type sequences conditioned on the unmasked residues, respectively. An obvious limitation of this kind of models is that they assume the same length of the mutant-type and wild-type sequences and thus cannot handle mutants resulted from indel (insertion/deletion) operations. In contrast, our formulation does not rely on the same-length assumption; instead, it sums over all residues (*L*) rather than only the mutated ones (*M*), making it suitable for mutations involving insertions or deletions. Moreover, the design leveraging unmasked predictions instead of masked ones allows other residues to account for mutation effects by aggregating information across the entire sequence too.

##### Motif-level Scoring Module

Motifs, which are consequential sequence patterns, play a crucial role in protein sequence modeling, widely used in bioinformatics research [64, 67]. In this work, we leverage convolutional neural networks (CNNs) to capture the local features of protein sequences. Specifically, we employ a convolutional module with max pooling and skipped connection to quantify the fitness from the local feature perspective. The score is computed as

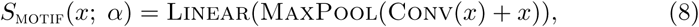

where *x* denotes the input sequence (of vectors) to the convolutional module, Conv and MaxPool denote the convolution operation and max pooling operation, respectively, and *α* are the parameters to be learnt in this module. The skipped connection is used to combine the output of the convolutional module with the input sequence, which helps to preserve the information from the original sequence. The score, *S*_motif_, is calculated by applying a linear transformation to the output of the convolutional module.

##### Sequence-level Scoring Module

It has been verified [21, 68] that the embedding of the [CLS] token in a natural language sentence is a good representation of the whole sentence. Therefore, in this work, the sequence-level scoring module in *µ*Former takes the embedding of the [CLS] token of a protein sequence as its representation. A Multi-Layer Perceptron (MLP) is employed to map the sequence representation to a fitness score, which is defined as

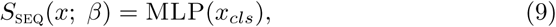

where *x_cls_* denotes the embedding of the [CLS] token generated by the pre-trained protein LM and *β* is the parameters to be learnt for this scoring module.

#### 4.1.3 Training

While the parameters *θ* of the protein LM is pre-trained using proteins without annotations, we need a set of proteins with measured fitness scores to train the newly introduced parameters (i.e., *α* and *β*) and to refine *θ*.

Let D denote a dataset consisting of pairs (*x, y*), where *x* represents a sequence of the wild-type or mutant protein, and *y* is the fitness score of the sequence. All the parameters are trained to minimize Mean Absolute Error (MAE):

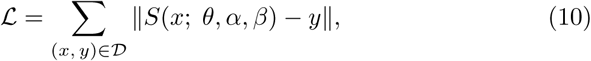

where *S*(*x*; *θ, α, β*) is the final fitness score obtained by summing three scoring modules.

To prevent overfitting the training data, which is likely derived from limited observations in biological experiments, the scoring modules are designed with significantly fewer parameters than the large, pre-trained encoder. Meanwhile, protein sequences typically consist of hundreds of amino acids, each with 20 possible alternatives. This necessitates a comparable number of parameters to effectively capture the information they contain.

#### 4.1.4 Implementation

The protein language models used in this study were pre-trained on the UR50 (release 2018 03) dataset. The base model was configured with a hidden size of 768, feed forward dimension of 3072, 12 encoder layers, and 12 attention heads. Additionally, a larger model was pre-trained with a hidden size of 1280 and 34 encoder layers. The base model was trained with a maximum length of 512, while the larger model was trained with a maximum length of 1024. Both models utilized non-learnable positional encoding. During pre-training, the base model used the Adam optimizer with parameters (0.9, 0.98), a peak learning rate of 0.0003, and a clip norm of 1.0. The learning rate was scheduled with a polynomial decay function, gradually decreasing after a warm-up period of 20, 000 steps. The larger model followed a similar hyperparameter configuration, with the peak learning rate set to 0.0001. Two models, *µ*Former-S and *µ*Former, are built on these two pre-trained models to assess the performance, with parameter sizes of 89 million and 670 million, respectively. *µ*Former-S and *µ*Former are trained using the same set of hyper-parameters across all datasets for a maximum of 300 epochs, with a hidden size of 256 and a motif scoring layer of 1. This indicates the potential for further performance improvement by tuning hyper-parameters for specific datasets.

#### 4.1.5 Datasets

**FLIP** [27]. The data is downloaded from Github: https://github.com/J-SNACKKB/FLIP/tree/main/splits/. We apply the original train/valid/test splits for evaluation, including the zero-shot models.

**ProteinGym** [5]. The data is called ProteinGym v0.1 and was downloaded from https://github.com/OATML-Markslab/ProteinGym on 12/30/2022, and this data version can be accessed via https://huggingface.co/datasets/OATML-Markslab/ProteinGym_v0.1 now. The substitution dataset is composed of approximately 1.5 million variants from 87 assays. Considering modeling efficiency, assays with target proteins longer than 1024 amino acids are removed, leaving 78 assays in the final analysis. The indel dataset is composed of approximately 300,000 mutants from 7 assays. Our experiments were conducted on ProteinGym v0.1 using a different data split scheme, where the data was randomly divided into training, validation, and test sets with a 7:1:2 ratio. We have made the dataset publicly available at https://doi.org/10.6084/m9.figshare.26892355. The performances of all trained models, including the zero-shot models, are reported on the fixed hold-out test set.

#### 4.1.6 Evaluation Protocol

For the supervised learning setting, we apply the original data split scheme for the FLIP benchmark, and evaluate the metrics fairly on the same hold-out test set. For the datasets from ProteinGym, we randomly select 10% of records from each dataset as a hold-out test set to evaluate all models rigorously including the zero-shot models. The performance numbers are reported on the fixed hold-out test set for a fair comparison. Spearman’s rank correlation scores are recorded for each dataset.

#### 4.1.7 Baselines

The multiple sequence alignments (MSAs) of the target wild-type sequences are prepared in advance using HHblits, by searching against the Uniclust30 database. For the ProteinGym datasets, the author provided MSAs are utilized. During our experiments, all models requiring evolutionary information features, including site-independent, EVE, EVmutation, DeepSequence, and ECNet, are trained on the same MSAs.

For the Tranception [5] model, we downloaded weights of the largest model (denoted as Tranception-L) from official website for zero-shot prediction. To generate zero-shot predictions using ESM-1v [24], we evaluated the ensemble results of 5 checkpoints downloaded from the official website. For the supervised setting of ESM-1v, we compared the results for the mean over subset, which was referred to as ‘mut mean’ in the FLIP benchmark [27]. This setting resulted in better overall performance than other settings evaluated on the benchmark. In this setting, fitness scores were calculated by regressions on sequence representations that were mean-pooled over the residues in the mutated region.

For supervised models, we use publicly available implementations and retrain them on the same data splits for each dataset. Specifically, for onehot embedding-based Ridge and CNN models, we adopt the architecture and hyperparameters from the FLIP codebase [27]. For ECNet [14], we reproduce the model using the open-source code provided by Luo et al. [14]. Since ECNet utilizes both MSAs and labeled data, we retrain it under the same settings using the prepared MSAs. Additionally, we fine-tune ESM-1b following the approach in [15]. We also include augmented Ridge models built on EVmutation and DeepSequence [15], evaluating both sampled and full-training data versions. For further comparison, we incorporate recent models, including ProteinNPT [32] and ConFit [33], along with its augmented variant leveraging DeepSequence.

### 4.2 Fitness Landscape Navigation by *µ*Search

To efficiently explore the vast space of mutations and identify high-functioning mutants in the protein landscape, we develop a reinforcement learning (RL) method called *µ*Search, which uses the *µ*Former model as a reward function. Similar to previous work [53, 70], we frame the protein engineering problem using a Markov Decision Process (MDP). In our MDP, the state represents the current mutant sequence, the action signifies a one-amino acid mutation, including the selection of mutation sites and types to mutate within the sequence, and the reward function is given by the *µ*Former model. With a protein engineering agent, during each episode of the MDP, the agent sequentially samples actions from a conditional distribution known as the policy (i.e., mutates a single-site residue) until it reaches a predetermined horizon. At the end of an episode, the agent produces a multi-point mutant from the wild-type sequence we are engineering. The objective of reinforcement learning is to optimize the selection policy for mutation sites and types for the desired properties through trial-and-error interactions with the reward function. This learning process alternates between two phases: mutation space exploration, and policy and value network updating. During the mutation space exploration phase, the agent navigates the extensive mutation space, leveraging a mutation site policy network and a mutation type policy network to generate potentially high-performing mutants. In the policy and value network updating stage, the agent uses the *µ*Former model to evaluate the newly generated mutants and employs update loss functions to improve the policy and value networks, aiming to propose mutants with higher fitness scores in subsequent iterations.

#### 4.2.1 PPO Algorithm with Mutation Site and Mutation Type Policy Networks

In *µ*Search’s MDP definition, each action corresponds to a single mutation which includes the selection of mutation sites and types within the sequence. Therefore, we have designed two distinct policy networks. The first, referred to as the mutation site policy network, uses both the original (wild-type) sequence and the current mutant sequence as inputs. These inputs are represented by a binary matrix of size 20 × 2*L*, where *L* denotes the length of the protein sequence and 20 means the number of types of amino acid. This network identifies the positions for modifications in the current mutant sequence. The second network, named the mutation type policy network, considers the wild-type sequence, current mutant sequence, and the selected mutation site, represented by a binary matrix of size 20 × 2*L* + *L*. This network determines the specific modifications to be made at the selected mutation site. The two networks work sequentially. First, the mutation site policy network identifies potential sites for mutation in the sequence. Then, the mutation type policy network selects the specific amino acid change to be made at the chosen site. We employ the Proximal Policy Optimization (PPO) algorithm [71] to optimize both policy networks, a method recently utilized to train large language models with human preference-based reward models [72, 73]. To reduce the variance of the policy gradient estimate, we incorporate two value networks that estimate the future rewards from a given state. The policy and value networks are updated by the following loss functions:

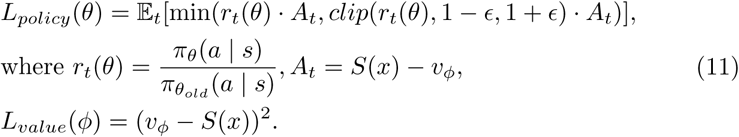

In this equation, *t* represents the step (timestep) in an episode, where each step corresponds to a single mutation being performed. The term *r_t_*(*θ*) is the probability ratio of the current policy *π*(*θ*) to the previous policy *π_old_*(*θ*), and *v_ϕ_* denotes the value function. *S*(*x*) is the *µ*Former score of the generated mutant *x*. The parameters *θ* and *ϕ* refer to the policy and value network parameters, respectively. Both the mutation site and mutation type policy networks use this loss function to update their model parameters.

The value function *v_ϕ_* serves as a baseline or critic, aiding in the computation of the advantage function *A_t_*. This function assesses the relative value of each action by calculating the difference between the actual return, *S*(*x*), and the expected return derived from the value function. The advantage function *A_t_* allows the policy network to differentiate superior actions from the rest, adjusting its probability correspondingly. The policy loss *L_policy_*(*θ*) is designed to increase the likelihood of actions with positive advantage functions while minimizing substantial deviations from the previous policy to prevent performance collapse. The hyperparameter *ɛ* controls the size of the trust region for each policy update.

#### 4.2.2 Incorporating Dirichlet Noise into Exploration

The mutation space of a protein sequence is vast, with high-profile mutants potentially spread widely within it. To navigate this space, we integrate Dirichlet exploration noise during the mutation generation phase, which helps prevent getting trapped in local optima. Specifically, the Dirichlet noise supplements the probabilities derived from the policy networks, resulting in *P* (*s, a*) = (1 − *ɛ*) · *π_θ_*(*a* | *s*) + *ɛ* · *η*, where *η* follows a *Dir*(0.03) distribution and *ɛ* is set to 0.25. This added noise ensures that all potential moves can be explored by the agent, while still prioritizing mutations with superior fitness scores from *µ*Former. During the updating phase, the injected Dirichlet noise may lead to an action with a low probability from the policy network *π_θ_*, potentially resulting in numerical optimization challenges when estimating the original importance weights, we introduce a ratio 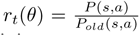 based on the mixed distribution, to facilitate a smoother training process.

#### 4.2.3 Searching with Multiple *µ*Former Models and Ranking based Selection for Experimental Validation

To identify TEM-1 variants with 2–3 amino acid mutations that exhibit enhanced activity against cefotaxime, *µ*Protein integrated the *µ*Former model with the *µ*Search algorithm. Six separate *µ*Search runs were performed using *µ*Former models trained with different random seeds. In each run, we set a threshold determined by the median predicted value of known ESBL mutants (as scored by the specific *µ*Former model used in that search). Variants exceeding this threshold were identified as high-potential candidates. For the TEM-1 system, a total of one million mutant sequences were computationally screened by the *µ*Former reward model across these six runs.

The results from the six independent searches were compiled into a comprehensive list of over 1,200 unique mutants. For each mutant in this list, we calculated its predicted scores across all six *µ*Former models and determined its relative rank within each search. The final ranking of the mutants was obtained by aggregating these relative ranks across the six searches. This automated ranking process was used to select the top 200 variants for experimental validation, balancing predictive confidence and wet-lab cost constraints. For the TEM-1 system, we conducted only one round of wet-lab experimental screening.

This approach ensures that the selection process is fully automated, free from manual curation, and leverages consensus predictions across multiple independent *µ*Former models for robust candidate prioritization.

### 4.3 *E. coli* growth assay

#### 4.3.1 library construction

The plasmid containing the wild-type TEM-1 sequence (pSkunk3-TEM-1) was generously provided by the Ostermeier lab. Amino acid substitutions proposed by *µ*Former were translated into corresponding DNA mutations, taking into account the codon preference in *E. coli*, and introduced into the plasmid by Suzhou Silicon BioScience. Engineered plasmids were verified by Sanger sequencing to confirm the presence of the intended mutations before being transformed into the SNO31 *E. coli* strain harboring pTS42, which was also provided by the Ostermeier lab. To facilitate downstream sequencing, a unique 6 bp strain-specific barcode was introduced into each TEM-1 mutant construct.

#### 4.3.2 Antibiotic Selection

Following established protocols from previous studies [45, 74, 75], bacterial cultures carrying various TEM-1 mutant plasmids were grown overnight to saturation at 37^◦^C, then diluted to an *OD*_600_ of 0.01. These cultures were seeded onto 180mm petri dishes containing LB media supplemented with either 0*µ*g/ml or 0.2*µ*g/ml cefotaxime sodium salt (TargetMol, CAS Number: 63527-52-6). In the initial round, around 50 different constructs were seeded onto the same plate, including the wild-type construct as a control. To ensure robustness, constructs were randomized across two different plates, each with distinct neighboring constructs. After 36-48 hours of incubation at 37^◦^C, bacterial colonies were harvested. The bacteria from each plate were collected into separate tubes for subsequent analysis and sequencing. The top 20 strains (described below) were selected for a second round of experiments where each variant, along with the wild-type, was individually seeded onto 90mm plates.

#### 4.3.3 Sequencing and analysis

Paired-end primers were designed to sequence the regions flanking the 6 bp barcode on the plasmid. A 3 bp sequencing barcode was incorporated into the sequencing primers to differentiate between plates. Barcoded PCR amplicons were prepared from each plate, pooled, and subjected to paired-end second-generation sequencing. Sequencing raw data was cleaned by removing adapters, trimming reads at the front and tail, and filtering out reads with *>* 40% unqualified reads (*< Q*15) using fastp (version 0.23.4). Paired reads were then merged based on at least 30bp of overlap. Sequence alignment was performed by minimap2 [76] (version 2.26) with the following parameter settings: “-a – sr -t 4 –cs -m20 –heap-sort=yes –secondary=no”. Aligned data were checked for the presence of the corresponding 6 bp barcode at the correct position. Only sequences with an exact match to the barcode were included in the downstream analysis. The relative abundance of mutant strains compared to the wild-type on each plate (log(FoldChange)) was used as a surrogate measure for the activity of TEM-1 mutants against cefotaxime.

The antibiotic selection, sequencing, and analysis were performed by Suzhou Silicon BioScience.

## Data Availability

The FLIP benchmark is publicly available at http://data.bioembeddings.com/public/FLIP/. The ProteinGym v0.1 data collection can be accessed at https://huggingface.co/datasets/OATML-Markslab/ProteinGym_v0.1. The dataset under our split scheme is available at https://doi.org/10.6084/m9.figshare.26892355. The list of extended-spectrum beta-lactamases can be found at http://bldb.eu/([44]). The fitness scores for beta-lactamase variants against cefotaxime, validated by our wet-lab experiments, are provided in the supplementary files.

## Code Availability

The code to reproduce the results of the paper is publicly available at the following URL: https://github.com/microsoft/Mu-Protein (https://www.doi.org/10.5281/zenodo.15836168). The data split scheme is available at https://doi.org/10.6084/m9.figshare.26892355.

## Acknowledgments

We gratefully acknowledge Dr. Ostermeier for generously providing detailed information on the plasmids pSkunk3-TEM-1 and pTS42, and for sharing the pTS42 plasmid itself. We also extend our thanks to Tianbo Peng for developing the demonstration webpage and Jingyun Bai for creating the graphic illustrations. Lastly, we appreciate Dr. Bonnie Kruft and Dr. Usman Munir for their invaluable support in program management and coordination.

## Authors’ contributions

Conceptualization, L.H., H.S., and P.D.; methodology and modeling, L.H., H.S., G.L., Z.Z., Y.J., F.J., and L.W.; data curation, L.H., H.S., and P.D.; result interpretation, P.D., H.L., L.H., and C.C.; writing-original draft, L.H., P.D., H.S., and G.L.; writing-review, H.L., T.Q., C.C.; supervision, T.L.. All authors read and approved the final manuscript.

## Ethics approval

Not applicable.

## Consent to participate

Not applicable.

---

**Extended Data Figure 1.**
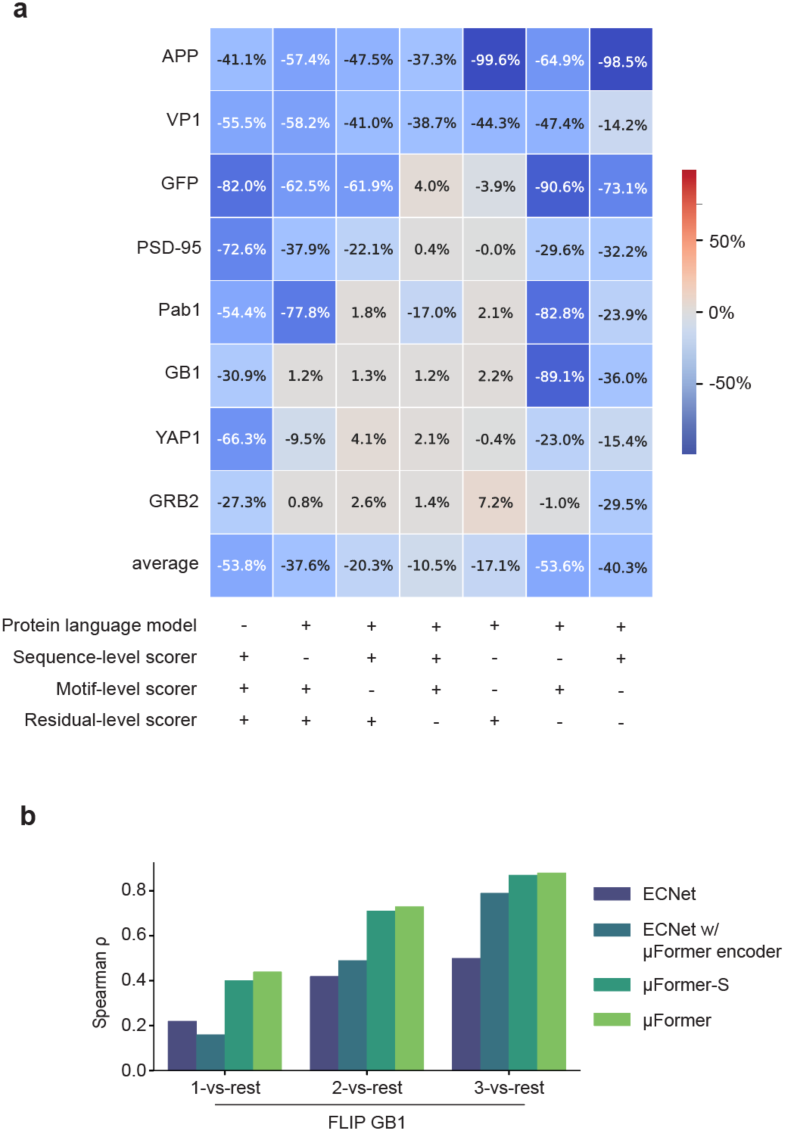
Ablation and analysis on μFormer components. **a)** Ablation study to evaluate the importance of each component in μFormer. The change in performance after removing various components from the model relative to a full model is shown. Negative numbers (blue) indicate a loss of performance and positive numbers (red) indicate an improvement in performance. The last row displays the average performance change over 9 proteins. The plus/minus signs at the bottom indicate the presence/removal of the corresponding component. **b)** Spearman ρ statistics on 3 FLIP GB1 datasets of μFormer, ECNet, and their variants. ECNet w/ μFormer encoder replaces the language model in ECNet with μFormer’s language model. μFormer-S (Methods) is a variation with a model size similar to ECNet. 1-vs-rest: a train-test split where single-point mutants are used for training, and multi-point mutants are reserved for testing. 2- vs-rest: a train-test split where single- and double-point mutants are used for training, and all higher-order mutants are reserved for testing. 3-vs-rest: a train-test split where single-, double-, and triple-point mutants are used for training, and all higher-order mutants are reserved for testing. See Supplementary Notes for details.

**Extended Data Figure 2.**
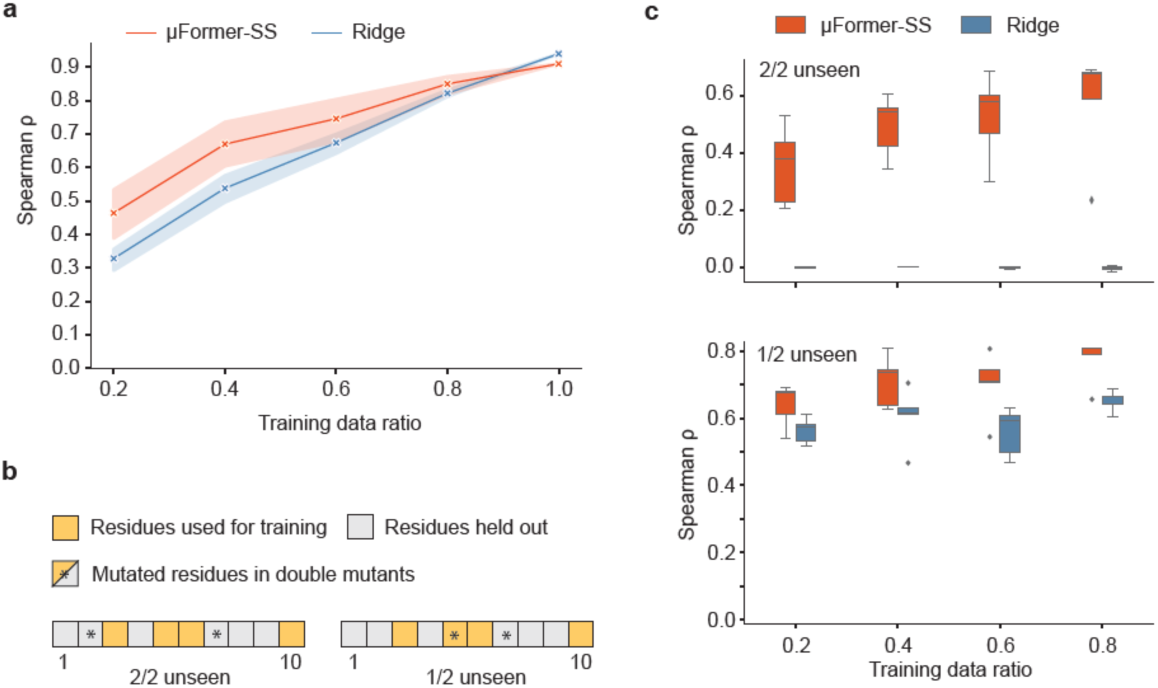
Analysis of μProtein. a) Performance of μFormer and Ridge on GB1 double mutants with varying training data size. Here, μFormer is a μFormer variation with a smaller supervised scorer module size (μFormer-SS). Training data ratio indicates the number of residues used for training versus the total number of amino acids in GB1. The training data size equals 209, 418, 627, 836, and 1045 for 20%, 40%, 60%, 80%, and 100%, respectively. All scores were evaluated on GB1 saturated double mutants (n=535,917). Center: mean. Error bands: standard deviation. Five experiments are performed for each setting with random selection on training data. b) Illustration of test data split, using a protein of 10 residues and the 40% setting as an example. 2/2 unseen: neither of the mutated residues in double mutants are seen by the model. 1/2 unseen: one and only one of the mutated residues in double mutants are seen by the model. c) Performance of μFormer and Ridge on different splits of GB1 double mutants. Training data split criteria are the same as in a). Center line, median; box limits, upper and lower quartiles; whiskers, 1.5x interquartile range; points, outliers. Five experiments are performed for each setting with random selection on training data.

**Extended Data Table 1:** Size of training (single-point mutants) and test (multi-point mutants) datasets in single-to-multi setting.

| DATASET | NO. OF TRAINING | NO. OF TEST |
| --- | --- | --- |
| GB1 | 1,045 | 535,917 |
| TEM-1 | 5,468 | 12,374 |
| Pab1 | 1,187 | 36,521 |
| PSD-95 | 1,280 | 5,696 |
| GRB2 | 1,034 | 62,332 |
| APP | 468 | 14,015 |
| GFP | 1,084 | 50,630 |
| YAP1 | 362 | 9,713 |
| VP1 | 532 | 41,796 |

