## Supplementary Information for "Accelerating protein engineering with fitness landscape modeling and reinforcement learning"

### Table of Contents

#### Contents

**1 Supplementary Notes** **2**
1.1 Ablation studies reveal the contribution of each component in $\mu$Former . . . 2
1.2 $\mu$Protein enables generalization to novel residues in target proteins 3
1.3 $\mu$Search efficiently navigate fitness landscapes of proteins (Extended) . . . 3

**2 Supplementary Figures** **6**

**3 Supplementary Tables** **18**

### 1 Supplementary Notes

#### 1.1 Ablation studies reveal the contribution of each component in $\mu$Former

To assess the contribution of each component in $\mu$Former, we conducted a series of ablation studies following a single-to-multi setting, where model is trained on single-point mutants only but required to predict fitness scores of multi-point mutants. We collected eight sets of DMS assays that include both single-point mutants and high-order mutants [1–8] and evaluated on each dataset independents.

As shown in Extended Data Fig.1a, omitting the language model results in the most significant drop in the performance of high-order mutational effect prediction, indicating the crucial role of pre-training in supervised tasks with limited labeled data. Meanwhile, scoring modules at different levels exhibit diverse roles for different target proteins, and combining the three modules results in the most robust performance. In most settings, disabling any one of the three scoring modules leads to a significant performance drop. However, for certain datasets, the model performs better when both the sequence-level and motif-level scorers are ablated compared to ablating either module individually. A plausible explanation is the relatively small size of the fine-tuning dataset. If these modules do not meaningfully contribute to fitness prediction, their inclusion may introduce noise or lead to overfitting, ultimately degrading performance. This also explains why the residue-level scorer does not exhibit the same behavior, as its parameters are derived from the pre-trained language model rather than being trained from scratch.

Moreover, to demonstrate that the multi-level scoring module design, rather than the model size or other factors, determines $\mu$Former's superior performance, we benchmarked $\mu$Former against three baselines on FLIP GB1 [8], a landscape focusing on four epistatic sites in GB1 protein with high mutational space coverage. $\mu$Former greatly outperforms ECNet, the learning-based method that also utilizes a language model, in all three settings provided by the benchmark dataset (Extended Data Fig. 1b). Additionally, replacing the language model in ECNet with $\mu$Former's language model (ECNet w/ $\mu$Former encoder) does not substantially improve performance, and a $\mu$Former variation with a model size similar to ECNet ($\mu$Former-S) exhibits a more similar performance to $\mu$Former (Extended Data Fig. 1b).

To evaluate the effect of our pre-trained protein language model in $\mu$Former, we conducted a series of comparative experiments. In addition to the PMLM model, we included the ESM-1b model for comparison, as both are pre-trained on the same sequence data. For a fair evaluation, we used identical scorer settings for both models. The experiments were carried out on the GB1 and AAV datasets, with the latter also sourced from the FLIP benchmark [8], following its prescribed data split schemes. Specifically, in the $n$-vs-rest setting, sequences with fewer than $n$ mutations are used for training and validation, while the remaining sequences are reserved for testing. The results, summarized in Supplementary Data Table 1: the performance gap for GB1 (2-vs-rest) and AAV (1-vs-rest) is substantial, likely due to the limited size of the datasets and the presence of low-order mutations used for fine-tuning. In contrast, for GB1 (3-vs-rest), where the training data is more abundant, the encoder differences become less pronounced.

#### 1.2 $\mu$Protein enables generalization to novel residues in target proteins

Considering that saturated single-point mutagenesis assays is often impractical to probe protein fitness landscapes at large scales, we next investigated whether $\mu$Former could generalize to residues unseen by the model. We used a saturated mutational effect assay of GB1 single and double mutants [6] and followed the single-to-multi setting aforementioned. We randomly selected 20%, 40%, 60%, and 80% of GB1 protein residues and trained the models with single mutant data exclusively covering these residues. For $\mu$Former, we decreased the size of the supervised scoring modules to avoid overfitting (denoted as $\mu$Former-SS). Subsequently, all prediction models were evaluated with saturated double mutants. We compared $\mu$Former to Ridge, a model demonstrating favorable performance in general evaluation. With five random repeats for each selection ratio, we found that $\mu$Former exhibited higher data efficiency and less sensitivity to the size of training data, achieving an average Spearman $\rho$ of 0.46 when only 20% of residues were used for training (Extended Fig. 2a). Furthermore, for each ratio and each repeat, we stratified the test dataset based on whether the two mutation sites were present in the training data (Extended Fig. 2b). Under the 20% setting, $\mu$Former reached an average Spearman $\rho$ of 0.36 when neither of the residues were seen (2/2 unseen) and 0.64 when one of the residues was seen (1/2 unseen) (Extended Fig. 2c). These results collectively indicate the data efficiency and generalization ability of $\mu$Former, leading to the conclusion that $\mu$Former can predict protein fitness and guide protein sequence design even with very limited data on the target protein.

#### 1.3 $\mu$Search efficiently navigate fitness landscapes of proteins (Extended)

In this set of experiments, we compared $\mu$Search against the following exploration algorithms:

- **AdaLead** [9]: an adaptive greedy search algorithm for sequence design;
- **DyNA-PPO** [10]: a reinforcement learning approach using proximal policy optimization for *de novo* biological sequence generation;
- **CbAS** [11]: a conditional sampling method for latent space optimization;
- **CMA-ES**: an evolutionary strategy with covariance matrix adaptation;
- **Bayesian Optimization**: a probabilistic model-based optimization framework;
- **PEX** [12]: a proximal exploration algorithm for local fitness landscape search;

- **GWG** [13]: a gradient-based Gibbs sampling method;
- **EvoPlay** [14]: a self-play reinforcement learning algorithm for protein engineering.

Additionally, we compared $\mu$Search with a Random Selection baseline, which uniformly selects mutation positions and types at random.

For the multi-round simulated experimental setup leveraging the FLEXS open-source simulation system [9], we assume a ground-truth oracle $\phi : x \to y$ that represents an expensive-to-query, ground-truth fitness function, as well as a local approximate oracle model $\hat{\phi}' : x \to y$ trained on samples $\mathcal{D}$ from the oracle $\phi$. The primary focus of this study is the performance of the exploration algorithm $\mathcal{E}$.

During each round, all exploration algorithms propose a batch of mutant sequences, constrained by a fixed budget of modeling calls to the local approximate model $\hat{\phi}'$. The proposed mutants are subsequently evaluated using the ground-truth oracle $\phi$, and the resulting evaluation scores are used to update the local approximate model $\hat{\phi}'$ for the next round. The cumulative maximum ground-truth fitness scores of all sequences proposed by each algorithm are plotted on the $y$-axis in Figs. 5a-c and Supplementary Data Fig. 6.

Notably, the same local and ground-truth models were consistently utilized across all exploration algorithms for five diverse fitness landscape exploration tasks provided by the FLEXS library (Supplementary Data Table 2):

- **TF-Binding Landscape** [15]: This landscape involves DNA 8-mers with fully characterized binding affinities. The sequence space ($4^8 = 65,636$) is exhaustive and the fitness landscape is governed by specific sequence motifs. This dataset is computationally simple and well-suited for baseline comparisons.
- **RNA Landscape** [16]: Simulated RNA binding landscapes generated using the ViennaRNA package, with sequence lengths limited to <200 nucleotides to ensure simulator accuracy. Fitness is determined by thermodynamic stability.
- **AAV Landscape** [17]: This landscape models the fitness landscape of AAV2 capsid protein using a Rough Mt. Fuji (RMF) additive model with Gaussian noise. The fitness landscape has a single global peak with predictable local structure and limited ruggedness. Sequence lengths range from short regions to full protein sequences.
- **GFP Landscape** [1]: Derived from the TAPE benchmark, this landscape involves optimizing fluorescence for 52,000 GFP variants of 238 amino acids.
- **Rosetta Landscape** [18]: This landscape uses PyRosetta for protein folding energy optimization (e.g., 3MSI antifreeze protein).

To evaluate the computational efficiency of $\mu$Search, we compared $\mu$Search to a random sampling baseline on exploring the TEM-1 landscape using the same $\mu$Former model as the oracle, consistent with the single-round setting in the manuscript. For each method, we measured the number of $\mu$Former model calls required to discover mutant sequences that surpass specified fitness thresholds. These model calls serve as a proxy for computational cost. We selected threshold values that appear in both methods to ensure a fair comparison. This setup allows for a direct comparison of the efficiency, where $M$ and $N$ denote the number of model calls made by random sampling and $\mu$Search, respectively. The results (Supplementary Data Table 3) demonstrate that $\mu$Search requires significantly fewer model calls to reach the same fitness level, especially for higher thresholds, indicating superior computational efficiency.

#### 2 Supplementary Figures

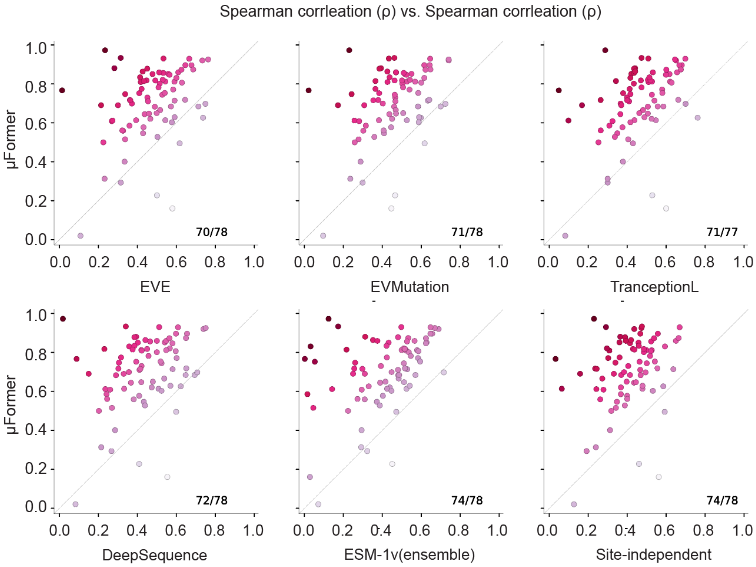


**Supplementary Data Figure 1. Pairwise comparison between μFormer and other approaches.** Each data point represents one ProteinGym dataset.

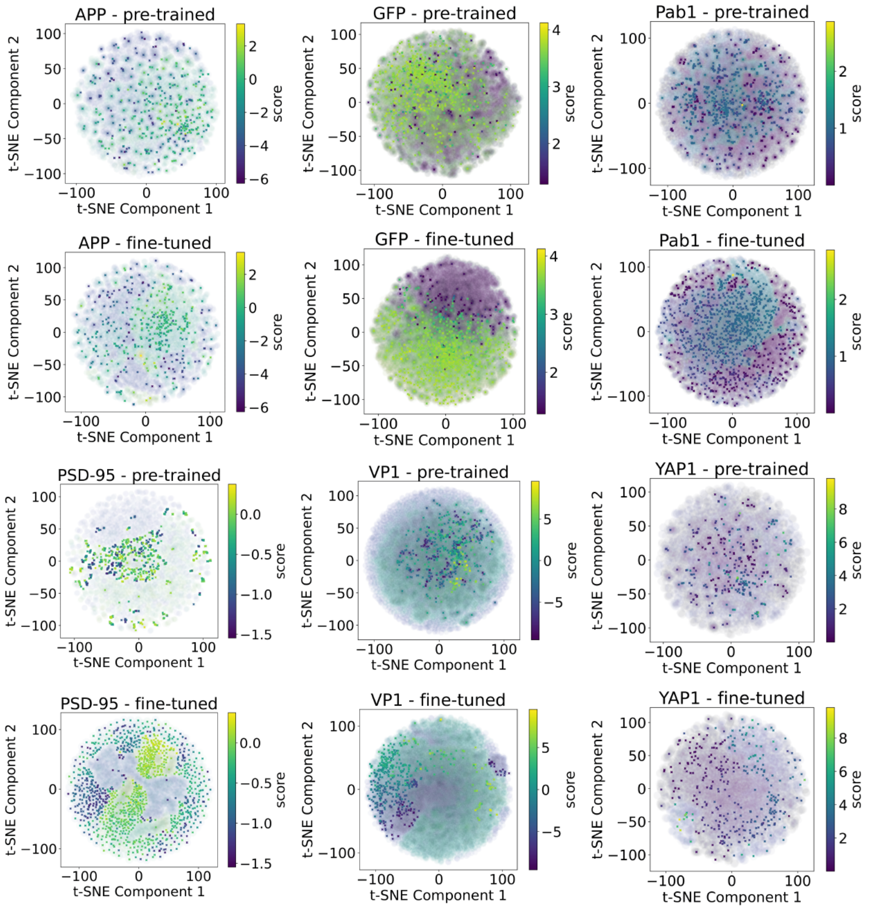


**Supplementary Data Figure 2. Visualization of embeddings on single mutants and high-order mutants (muti-point mutants) using t-SNE.** First row: Illustration of embeddings on APP, GFP, and Pab1 extracted from pre-trained μFormer model. Second row: Illustration of embeddings on APP, GFP, and Pab1 extracted from fine-tuned μFormer model. Third row: Illustration of embeddings on PSD-95, VP1, and YAP1 extracted from pre-trained μFormer model. Fourth row: Illustration of embeddings on PSD-95, VP1, and YAP1 extracted from fine-tuned μFormer model. The color scheme illustrates the fitness scores, with lighter shades indicating higher scores. Blurred background markers represent high-order mutants, while clear foreground markers denote single mutants.

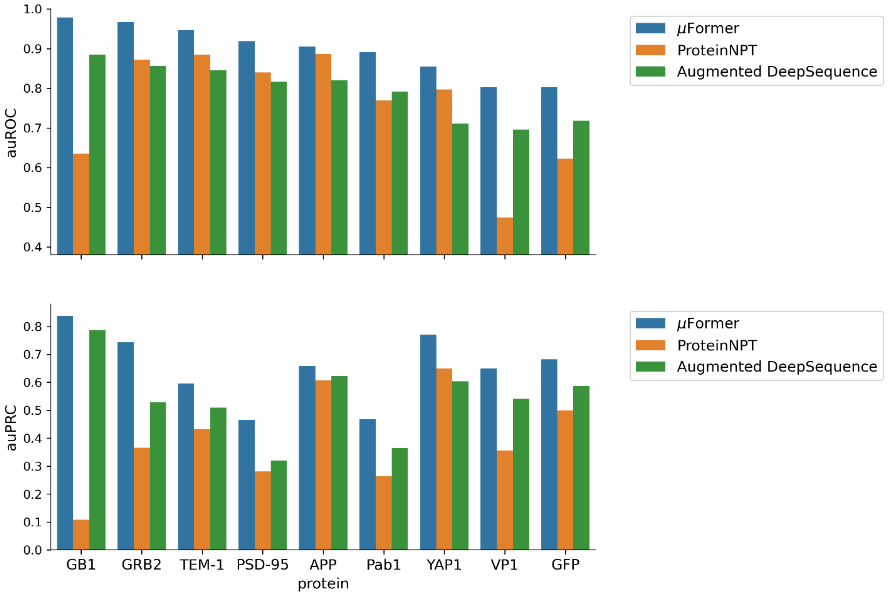


**Supplementary Data Figure 3. μFormer accurately distinguishes gain-of-function mutations from loss-of-function mutations.** Upper: the area under the receiver operating characteristic (auROC) score for different models on various proteins. Bottom: the area under precision-recall curve (auPRC) score for different models on various proteins.

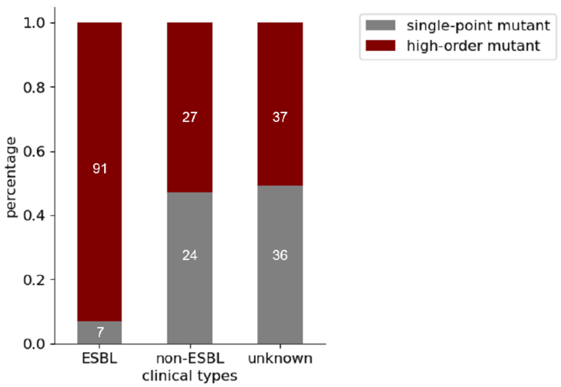


**Supplementary Data Figure 4. Percentage of single-point mutants and high-order mutants in each clinical type of TEM-1.** The count number for each group is marked on the plot.

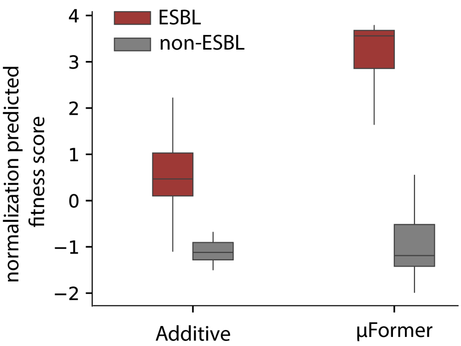


**Supplementary Data Figure 5. Normalized predicted fitness scores for the additive baseline and μFormer.** Z-score normalization was applied to ensure the predicted fitness scores from both models were on the same scale and distribution for comparison. For the additive baseline, fitness effects of all single mutants were first predicted using μFormer. The predicted effects were then summed relative to the wild-type sequence for all mutations present in each multi-mutant sample in the test dataset. This approach allows for a direct comparison of μFormer and the additive baseline in distinguishing ESBLs from non-ESBLs. Red: Normalized predicted fitness scores for ESBLs. Gray: Normalized predicted fitness scores for non-ESBLs. Center line, median; box limits, upper and lower quartiles; whiskers, 1.5x interquartile range; points, outliers. n = 156.

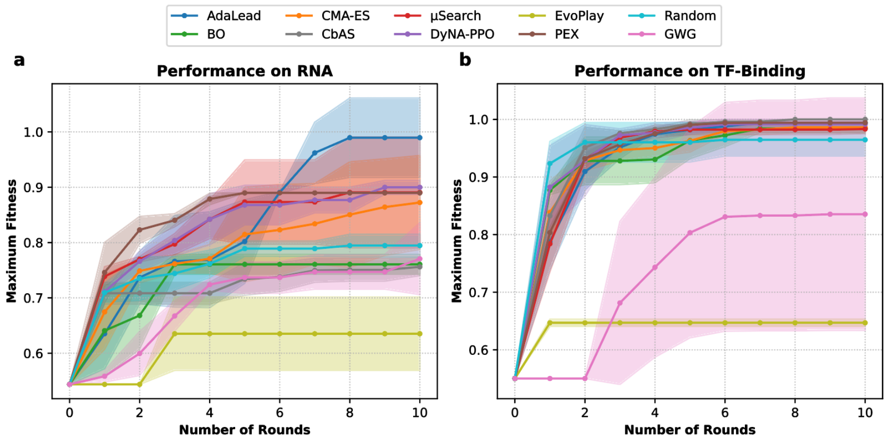


**Supplementary Data Figure 6. Quantitative comparison of μSearch with prevalent exploration methods across various fitness landscapes. a-b)** Compare μSearch with eight leading methods: AdaLead, DyNA-PPO, CbAS, CMA-ES, Bayesian Optimization, PEX, GWG, and EvoPlay, as well as a Random Selection baseline across simulated RNA binding landscape **(a)** and 8-mer DNA-Transcription factor binding (TF-binding) landscape **(b)**. Center: mean. Error band: standard deviation.

**a**

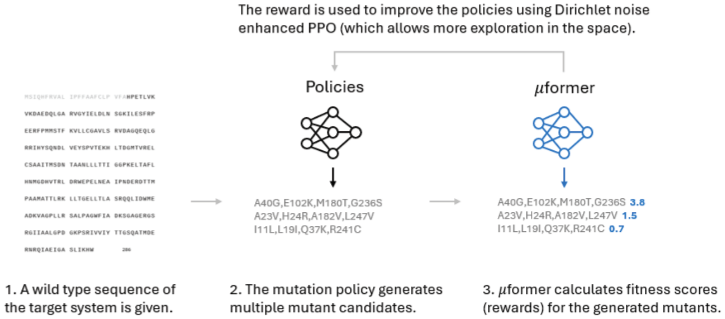


**Supplementary Data Figure 7. Reinforcement learning supports efficient and comprehensive exploration of the vast protein mutant space. a)** Diagram of reinforcement learning-based search pipeline.

**a)**

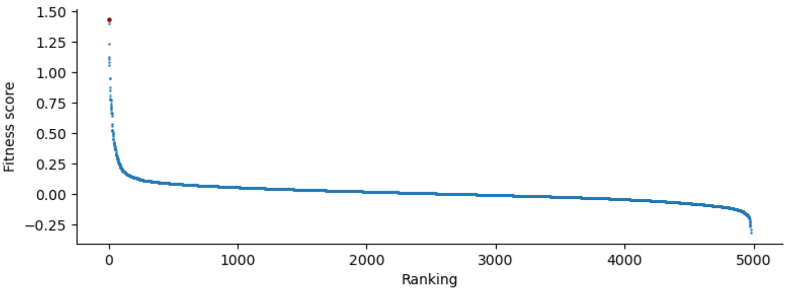


**b)**

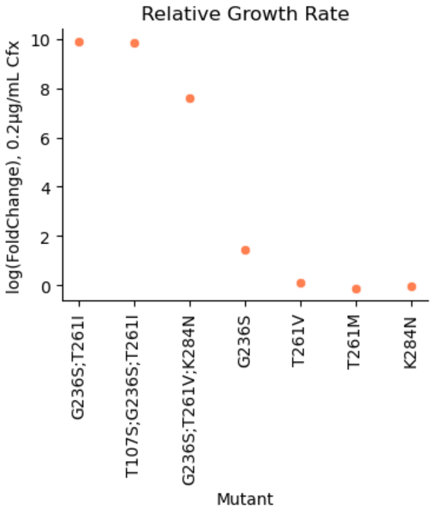


**Supplementary Data Figure 8. Analysis on TEM-1 mutants. a)** Ranking of the observed fitness scores for training data (Stiffler, 2015). G263S is highlighted in red. **b)** Fold change in growth of TEM-1 mutants compared to wild-type *E. coli* in the presence of 0.2 µg/ml cefotaxime.

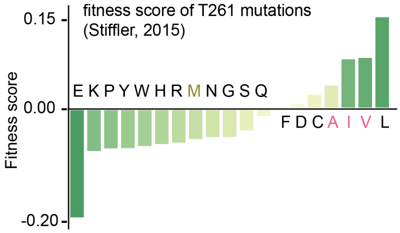


**Supplementary Data Figure 9. Fitness scores of various TEM-1 T261 mutations, as quantified by Stiffler et al. 2015.**

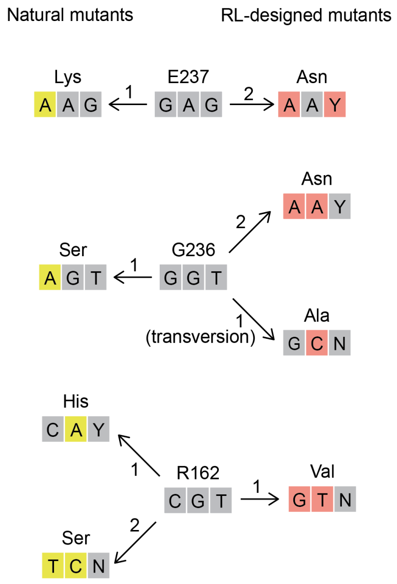


**Supplementary Data Figure 10. Nucleotide mutations in natural mutants (ESBL strains, left) and RL-designed mutans (right).** The numbers 1/2 represent the number of nucleotide mutations required for the amino acid substitution. Y denotes C/T, and N denotes A/T/C/G.

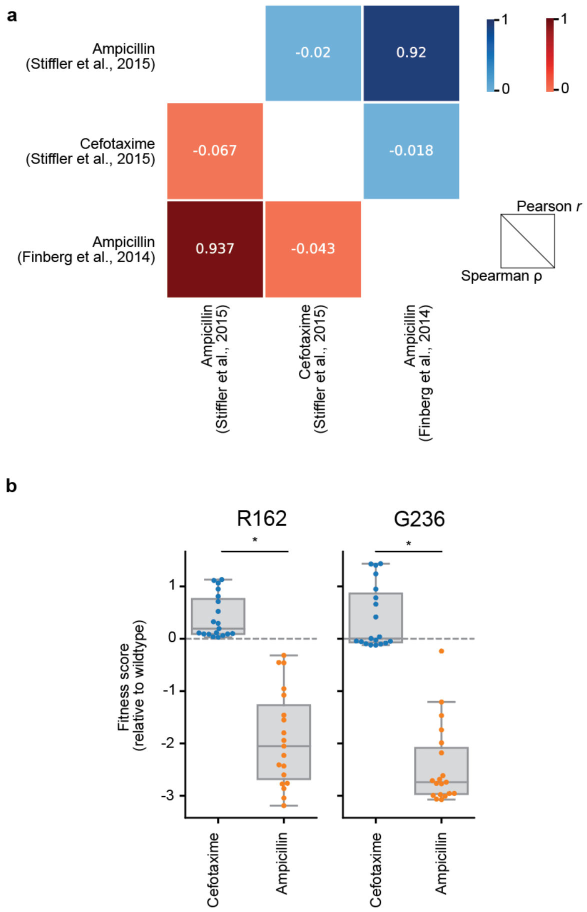


**Supplementary Data Figure 11. TEM-1 mutants' activity against ampicillin and cefotaxime are poorly correlated. a)** Spearman ρ and Pearson r representing the correlation between TEM-1's mutational effects on cefotaxime and ampicillin. The data on ampicillin was collected from 2 different studies. **b)** For residues R162 and G236, the same set of mutations exhibits contrasting effects on protein's activity against cefotaxime and ampicillin. Center line, median; box limits, upper and lower quartiles; whiskers, 1.5x interquartile range; points, outliers. P-values were calculated with two-sided Mann–Whitney U test. R162 p-value: 1.48e-07. G236 p-value: 1.48e-07. $n$ = 38.

### 3 Supplementary Tables

**Supplementary Data Table 1**: Ablation study for Protein LM Encoders

| SETTING | GB1 (2-VS-REST) | GB1 (3-VS-REST) | AAV (1-VS-REST) |
|---|---|---|---|
| $\mu$Former w/ ESM-1b | 0.61 | **0.88** | 0.57 |
| $\mu$Former w/ PMLM | **0.73** | **0.88** | **0.68** |

**Supplementary Data Table 2**: Benchmark landscape characteristics by domain and mutation space size

| LANDSCAPE | DOMAIN | MUTATION SPACE SIZE |
|---|---|---|
| TF-binding | DNA | $4^8 \approx 6.5 \times 10^5$ |
| RNA | RNA | $4^{100} \approx 1.6 \times 10^{60}$ |
| AAV | Protein | $20^{90} \approx 1.2 \times 10^{117}$ |
| GFP | Protein | $20^{238} \approx 4.4 \times 10^{309}$ |
| Rosetta | Protein | $20^{90} \approx 1.2 \times 10^{117}$ |

**Supplementary Data Table 3**: Computational efficiency comparison on the TEM-1 dataset

| FITNESS THRESHOLD | $M$ (RANDOM SAMPLING) | $N$ ($\mu$SEARCH) | $M$ / $N$ |
|---|---|---|---|
| -0.06 | 50 | 50 | 1.0 |
| -0.03 | 100 | 50 | 2.0 |
| 0.01 | 300 | 50 | 6.0 |
| 0.05 | 3,950 | 150 | 26.3 |
| 0.07 | 20,550 | 200 | 102.8 |
| 0.08 | >50,000 | 350 | >143.0 |
